# Improving diffusion-based protein backbone generation with global-geometry-aware latent encoding

**DOI:** 10.1101/2024.10.05.616664

**Authors:** Yuyang Zhang, Yuhang Liu, Zinnia Ma, Min Li, Chunfu Xu, Haipeng Gong

**Affiliations:** MOE Key Laboratory of Bioinformatics, School of Life Sciences, Tsinghua University, Beijing, 100084, China; Beijing Frontier Research Center for Biological Structure, Tsinghua University, Beijing, 100084, China; National Institute of Biological Sciences, Beijing, 102206, China; Tsinghua Institute of Multidisciplinary Biomedical Research, Tsinghua University, Beijing, 100084, China; Department of Bioengineering, University of California San Diego, La Jolla, CA 92093; China National Center for Protein Sciences, Beijing, 100084, China; X-ray Crystallography Facility, Technology Center for Protein Sciences, Tsinghua University, Beijing, 100084, China

**Keywords:** protein design, diffusion model, representation learning, controllable generation, mainly beta proteins

## Abstract

Recent breakthroughs in diffusion-based generative models have prompted *de novo* protein design, notably in generating diverse and realistic structures. Nevertheless, while existing models either excel at unconditional generation or employ residue-wise conditioning for topological control, explorations on a holistic, top-down approach to control the overall topological arrangements is still limited. In response, we introduce TopoDiff, a diffusion-based framework augmented by a structure encoder and a latent sampler. Our model can unsupervisedly learn a compact latent representation of protein global geometry, while simultaneously integrating a diffusion module to leverage this information for controlled structure generation. In benchmark against existing models, TopoDiff demonstrates comparable performance on established metrics and exhibits an improved coverage over the fold modes of natural proteins. Moreover, our method enables versatile control at the global-geometry level for structural generation, under the assistance of which we derived a number of novel folds of mainly-beta proteins with comprehensive experimental validation.

## Introduction

The *de novo* protein design is an intriguing and expanding field of research with the potential to venture into uncharted fold space, offering limitless opportunities for tailoring proteins to novel applications, including biomedical therapeutics^1–3^, catalytic enhancement^4^ and the development of innovative biological circuits^5,6^. Despite its vast potential, the *de novo* protein design has long been recognized as a challenging task, due to the highly structured nature of protein data and the stringent requirements on geometric restraints^7^.

The recent advance in the diffusion models significantly reshaped the field with its superior ability to generate novel, diverse and physically plausible structures. Though the pioneer efforts still relied on 1D or 2D protein representations^8–10^, subsequent works tended to leverage the success in protein structure prediction tasks^11,12^, building an equivariant network to directly learn the physical prior in the Cartesian space^13–17^.

Despite the encouraging progress, several issues remain to be addressed. While lacking systematical evaluation, there is evidence that some models, although trained on unbiased datasets such as the Protein Data Bank (PDB)^18^ or CATH^19,20^, struggle to generate protein backbones of certain fold classes^16^. This noteworthy fact becomes more evident when one reviews all experimentally validated proteins generated by structure-based diffusion models up to now, where the successful results are dominated by proteins of mainly-alpha or alpha & beta classes. Furthermore, we note that the current widely used metrics, namely the designability, novelty and diversity, provide no indications of the extent to which the natural protein space has been covered. This deficit further hinders the understanding and solving of the issues.

To improve the coverage of generated samples over specific protein folds, previous works have used residue-level 1D or 2D fold conditioning along with additional fine-tuning to generate immunoglobin domains with varied loop regions^21^, or applied classifier guidance by training classifiers on specific protein classes^22^. While these established methods can enhance coverage for particular groups of proteins, they are feasible only when such groups are clearly and distinctly defined and contain a sufficient number of training samples, which allows the application of its finer-grained topology as conditioning, the training of a robust classifier to guide the gradient, or even model fine-tuning on specific data subsets for additional refinement. However, due to the limited and unbalanced amount of available annotations as well as the discreteness and subjectivity in annotation assignment^23,24^, it is often impractical to apply the same strategy to cover all fold modes in the training set in an unbiased manner and to expand the protein fold space simultaneously.

In this work, we focus on an important and generic unsupervised problem setting. Given an arbitrary dataset, how can we train the diffusion model to capture the underlying data distribution without the explicit requirement of annotations or prior understanding of the dataset? Furthermore, can we leverage the power of state-of-the-art generative model design to learn interpretable latent encoding so as to deepen our understanding? Here, we propose the framework of TopoDiff as a solution. First, we concurrently trained the diffusion-based structure generative model with a structure encoder of the encoder-decoder architecture, thereby enabling the learning of a compact and fix-sized, continuous latent space that encodes the high-level global geometry of proteins as well as the simultaneous derivation of a generative module that operates at the residue level by conditioning on given latent information. The bottleneck design and distribution regularization jointly endow the encoder an inductive bias to capture a number of crucial degrees of freedom that allow the reasonable partitioning of the samples in the dataset, therefore presenting an invaluable, semantically meaningful latent space to facilitate not only the understanding of the dataset but also the controllable generation of new samples. Next, we trained a simple latent diffusion model to sample from the learned latent distribution unbiasedly and used the sampled latent to guide protein structure sampling, a scheme that effectively enhances the coverage of the fold modes in the dataset and opens up a new dimension for controllable generation at the same time. We then defined a coverage metric and conducted systematic evaluations on TopoDiff and other state-of-the-art models using this newly proposed metric and established ones including designability, novelty and diversity. Finally, we performed biological experiments to demonstrate the effectiveness of our model in improving the sampling of proteins of mainly beta topologies, a large class that is prevalent in nature but remains underrepresented in prior *de novo* protein design attempts. Specifically, out of the 21 candidates sampled *in silico* with novel mainly-beta topologies across a wide range of lengths, 9 proteins show expression in *E. coli*, among which 4 were successfully purified to obtain their monomeric forms. The circular dichroism (CD) spectra of these four proteins are consistent with the formation of predominantly beta-strand secondary structures. We further determined the high-resolution structure for one of the designs via X-ray crystallography and confirmed that the designed structural model matches the experimentally resolved structure with an high accuracy (RMSD of 1.31 Å).

## Results

### Overview of TopoDiff

The overall framework of TopoDiff is illustrated in Figure 1. The global structural properties of a protein, such as the shape, fold and topology, are closely related to its function^25,26^ and dynamics^27,28^. Similarly, when investigating protein structures, researchers always have a clear heuristic to first understand its global arrangement, followed by detailed examinations of the localized functional sites. Conversely, albeit powerful, classic diffusion models confine the dimensionality the same as the input, frustrating efforts to learn meaningful, compressed latent representations for the global geometry of proteins^29^.

**Figure 1.**
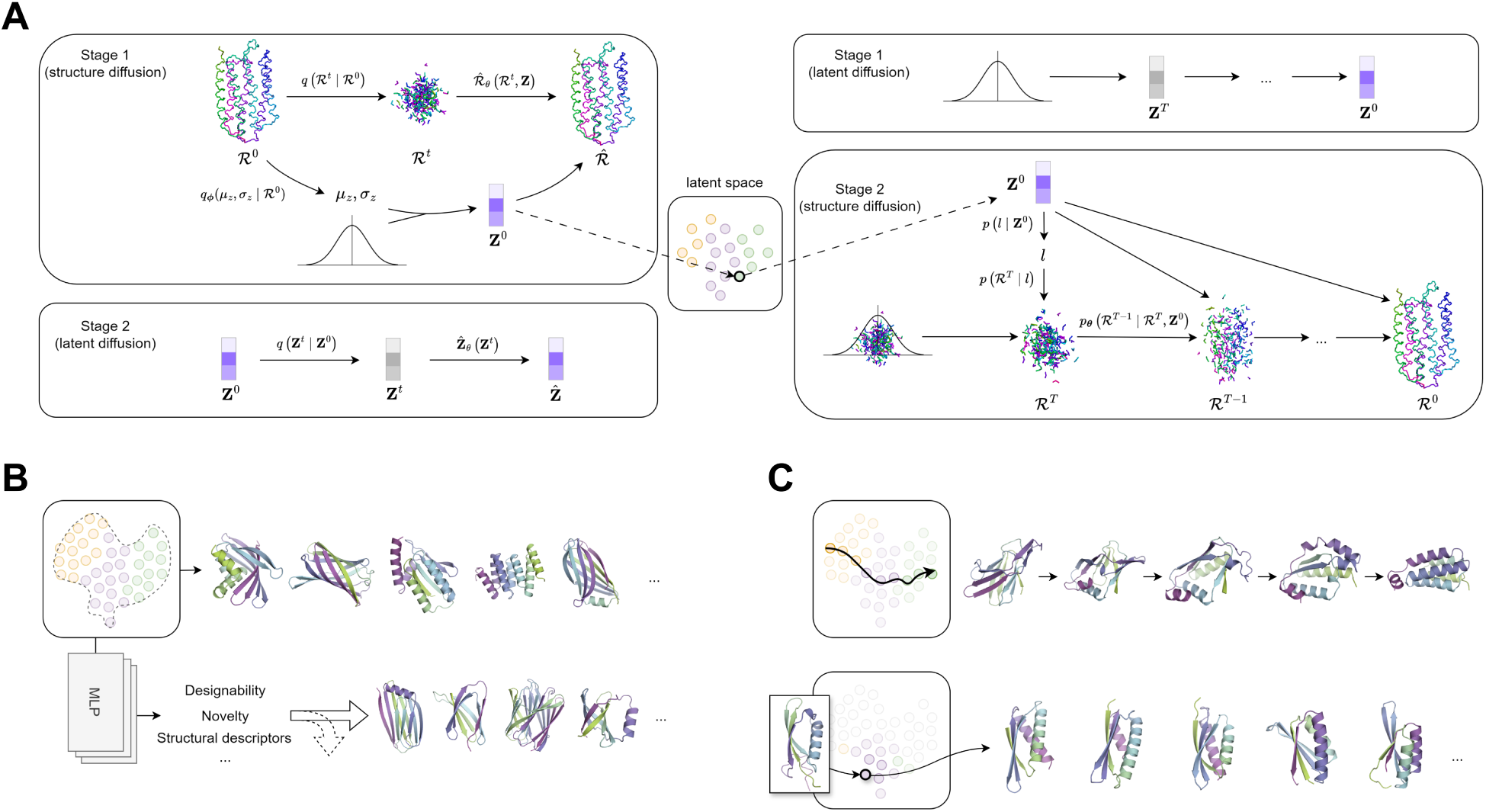
Overview of the Work. **(A)** The overall framework of TopoDiff. The training (left) and sampling (right) processes both consist of two stages. In Stage 1 of the training process, each protein structure is converted by the encoder module into a fixed-size, low-dimensional latent code **z** that captures the global geometry, whereas the perturbed structure *R^t^* is sampled following the designed forward marginal distribution. The diffusion decoder is trained to predict the ground truth structure *R*^0^ from *R^t^* and **z**. In Stage 2, the training focuses on learning the diffusion process in the latent representation space. During inference, a latent code is first sampled from the latent diffusion module, and then the structure decoder is employed to generate structures conditioned on the sampled latent. **(B)** Through unbiasedly sampling in the compact, continuous latent space, TopoDiff learns to generate samples with enhanced coverage of the natural fold space (upper row). Otherwise, TopoDiff could be targeted to produce structures with desired properties by reweighted latent sampling under the assistance of latent-based property predictors (lower row). The example here shows a particular application for generating novel, designable mainly-beta proteins. **(C)** TopoDiff is capable of generating structures from a local distribution in the latent space. The two exemplar applications are the simulation of gradual transition between two structures through latent-based linear interpolation (upper row) and the creation of new structures with similar global geometry to a given reference through latent perturbation (lower row).

Holding this notion, the core focus of our framework is on the establishment, investigation and utilization of a fixed-size, low-dimensional latent representation that encodes the global structural characteristics of a protein, building upon the latest advance of diffusion-based models. To achieve this aim, both the training and sampling processes are divided into two stages. In Stage 1 of the training process (Figure 1A, left), we adopt a diffusion-VAE formulation to harness the superior generative capability of diffusion models alongside the representational power of Variational AutoEncoders (VAE), thereby combining the inherent strengths of both architectures in a unified training process. The resulting structure encoder and conditional diffusion decoder, which share a common latent space, are the essential components of our framework. In Stage 2, a latent diffusion module is trained to model the latent distribution, enabling unbiased sampling from this otherwise intractable space. During the sampling phase (Figure 1A, right), the latent diffusion model first samples from the latent representation space, and a structure diffusion process is then performed for protein structure generation in the Cartesian space by conditioning on the sampled latent.

These steps describe the fundamental training-sampling loop of TopoDiff, whereas examples of new sampling schemes enabled by the TopoDiff framework are shown in Figure 1B&C. In the subsequent sections, we seek to progressively explore the unique characteristics of TopoDiff, focusing on its interpretability, generation quality and controllability.

### TopoDiff learns a compact and interpretable representation of the fold space

To gain an understanding of the latent space learned by the model, we first encoded all structures in our CATH-60 training dataset into vectors in the 32-dimensional latent space and then applied the t-SNE with the L2 distance metric^30^ for dimension reduction. As shown in Figure 2B, the latent codes collectively form a compact and continuous manifold. Notably, even though no structure annotations were used in our training process, the clustered structures perfectly coincide with the human curation of CATH class annotations, with each class clearly separable from the others even in the 2-dimensional reduced space. Moreover, we find that each CATH Architecture cluster indeed exhibits a unique spatial distribution (Figure 2C and Supplementary Figure S2). Many other intrinsic attributes of the proteins, such as the secondary structure proportions, chain length and radius of gyration, also exhibit structured global or local distribution patterns within the manifold (Supplementary Figure S1). These observations support that the model learns to partition over the training dataset in an unsupervised and highly interpretable manner.

**Figure 2.**
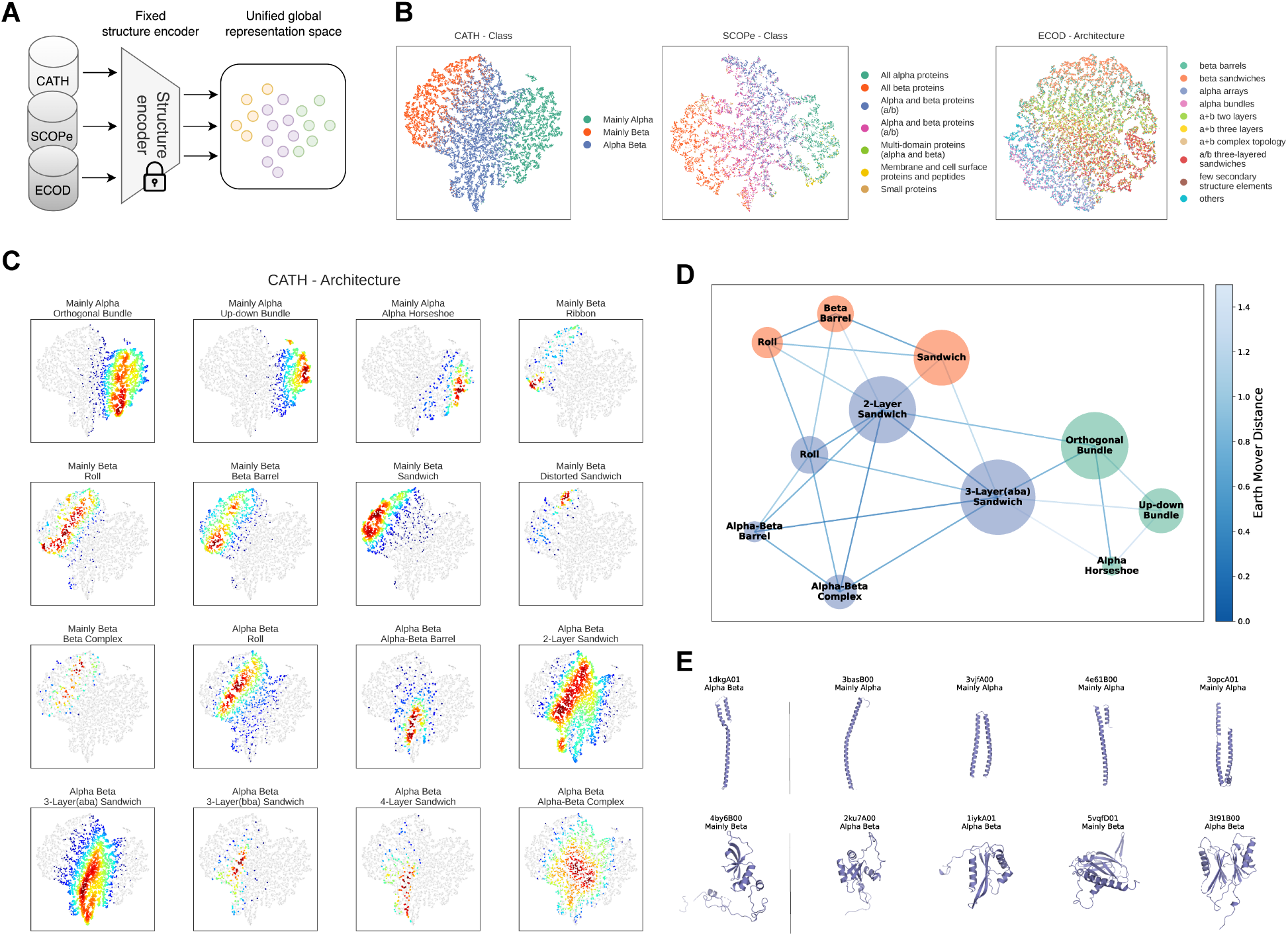
Analysis of TopoDiff’s Learned Latent Representations. **(A)** Once trained, the fixed structure encoder maps structures from different sources into a unified global representation space. **(B)** Visualization of the latent space colored by top-level classification hierarchies: CATH (Class), SCOPe (Class), and ECOD (Architecture). Each point represents a structure, colored according to its annotation. **(C)** Kernel density estimation of specific CATH architectures within the latent space. Each subplot shows the density of structures within a particular architecture, illustrating its distribution across the latent space. **(D)** Graph representation of several major CATH architectures. The nodes represent architectures, while the edges represent their relationships in the latent space, with the thicker and darker lines indicating the smaller Wasserstein distances. **(E)** Representative protein structures identified as outliers in their respective categories, along with their closest neighbors in the latent space. Each structure is annotated with its category (Class hierarchy or Architecture hierarchy) to explore the discrepancies between latent space clustering and database annotations.

Next, we tested the generalizability of this encoding method to unseen data. Specifically, we used the very same encoder trained on CATH to investigate two other structural classification datasets, SCOPe^31,32^ and ECOD^33^ (Figure 2A). Notably, although the three datasets all classify natural proteins at the domain level in a similar discrete and hierarchical manner, they adopt completely different classification schemes (in terms of methodology and underlying assumption)^34,35^, which leads to their evident distinctions in structural coverage and domain boundary definitions^34,36,37^. Interestingly, the latent manifolds produced by the encoder align well with the top-level annotations of the hierarchies in both SCOPe and ECOD datasets (Figure 2B and Supplementary Figures S3&S4). In line with past analyses^34,36,37^, the distinctive hierarchy organizations of different classification systems may essentially result from different domain partitioning and class discretization within a common structure space, for which we have sought to learn a continuous representation in an unsupervised manner and thus bypassed the annotation inconsistencies among datasets.

Moreover, the learned continuous latent space may supplement the information lost during the discretization of the natural fold space in traditional classification systems^20,31–33^. We take three node categories at the Architecture level of CATH hierarchy, Roll (CATH ID 2.30), Roll (CATH ID 3.10) and 3-Layer(aba) Sandwich (CATH ID 3.40), as an example to show the information loss in the CATH system. Based on the hierarchical classification, Roll (2.30) is equally distant to Roll (3.10) and 3-Layer(aba) Sandwich (3.40), two nodes with the sibling parents, although the former two categories clearly possess a stronger similarity indicated both by their names and by the spatial arrangement of beta-strand secondary structures. Here, we mapped CATH structures as points in the latent space and then computed the Wasserstein distance between the two sets of points (Equation 5) to quantify the relationship between a pair of CATH architectures. By this means, we constructed a graph diagram for a few major CATH architectures (Figure 2D and Supplementary Figure S5). Clearly, architectures from the same class hierarchy are generally closer to each other and the three CATH classes aggregate into distinctive clusters. In addition to these qualitative agreements, some continuous relationships between architectures emerge beyond the database annotations. For example, the Roll (3.10) architecture from the Alpha-Beta class exhibits the shortest distance to Roll (2.30) among all architectures in the Alpha-Beta and Mainly-alpha classes. Similarly, the 3-Layer(aba) Sandwich (3.40) is arguably closer to the Mainly-alpha class than its sibling architectures, whereas the 2-Layer Sandwich (3.30) presents similar distances from the Mainly-beta and Mainly-alpha classes. These observations are roughly consistent with the previous works^23,38^. Hence, the discretization of fold space by categorical classification inevitably leads to information loss, and by learning a generic representation from the ground up, our approach reveals the continuous relationships that have been overlooked.

On the other hand, at different hierarchical levels of the CATH system, we occasionally observed outliers whose latent representations are distantly located from the rest structures of the same CATH annotation. We conducted further filtering by only preserving the ones with non-uniform CATH annotations in their latent-space neighborhoods. Figure 2E shows two exemplar outlier structures along with their closest neighbors in the latent space and their corresponding annotations. The structural resemblance between the identified structures and their neighbors clearly indicates the inconsistency of CATH annotation assignment, which prioritizes evolutionary proximity over structural similarity when assigning new structures to annotated categories^34,39^. More cases could be found in Supplementary Figures S6&S7. In general, the identified outliers always present structural similarity to their latent-space neighbors to a considerable extent, despite the discrepancy in the CATH annotations. Deterministic categorical assignment is highly challenging for these structures due to the presence of structural heterogeneity within the CATH annotation categories, which is unable to be accounted for by discrete categorical metrics but could be adequately described using a continuous latent representation.

In summary, all findings in this section uniformly suggest that the encoder in TopoDiff learns a generic representation of the global structural geometry of natural proteins, offering an alternative continuous formulation that eliminates the need for discretizing and annotating individual structures. This encoding may provide assistance in the interpretation of the existing structure space and in guiding the diffusion-based protein structure generation as informative conditions.

### Benchmark testing on unconditional sampling

Here, we seek to evaluate the performance of TopoDiff in the unconditional sampling of new structures. As detailed in the “Methods” section, unconditional sampling of the entire space is achieved using ancestral sampling, where a latent code is sampled first through the latent sampler, followed by the structure sampling conditioned on that code. We compare with a few state-of-the-art diffusion-based generative models, including Genie^16^, FrameDiff^15^, Chroma^22^ and RFDiffusion^17^. To ensure a comprehensive and robust evaluation across a broad spectrum of protein sizes, for each model under evaluation, we randomly generated 500 structure samples at each fixed length among {50, 75, 100, 125, 150, 175, 200, 225, 250}, a series uniformly spanning the range of samples in our training data. Based on our experience, such an evaluation scheme can detect the length-dependent behavior in a model, which might be overlooked otherwise.

We notice that established metrics are dedicated to quantifying either the per-sample quality (*e.g.*, designability and novelty) or the within-sample diversity (*e.g.*, diversity) but provide no information of how much the known fold space is covered by the generated samples, an indicator that quantitatively describes the model capacity in unbiased sampling among existing data. Evidence from other domains has shown that the lack of considering this metric will introduce selection bias towards models that sacrifice variation in favor of high-quality samples from a truncated subset of the sampling space^40^, leading to substantial disruption to the model applicability. Indeed, over the past decade, the *de novo* protein design has been largely confined to alpha helix bundles and alpha-beta sandwiches^41,42^, and diffusion-based models have not provided an immediate remedy to this biased trend, as the experimental validations and applications still predominantly focus on these architectures^17,22,43^. To address this problem, we adopted a coverage metric^44^ to quantify the extent to which natural protein folds are covered by the samples produced by a generative model. Specifically, we used the CATH-40 non-redundant dataset as a representation of natural proteins and measured the fraction of proteins whose KNN (*i.e. K*-nearest neighbors) neighborhood contains at least one generated sample (Figure 3A). The definition and implementation details are provided in the “Methods” section.

**Figure 3.**
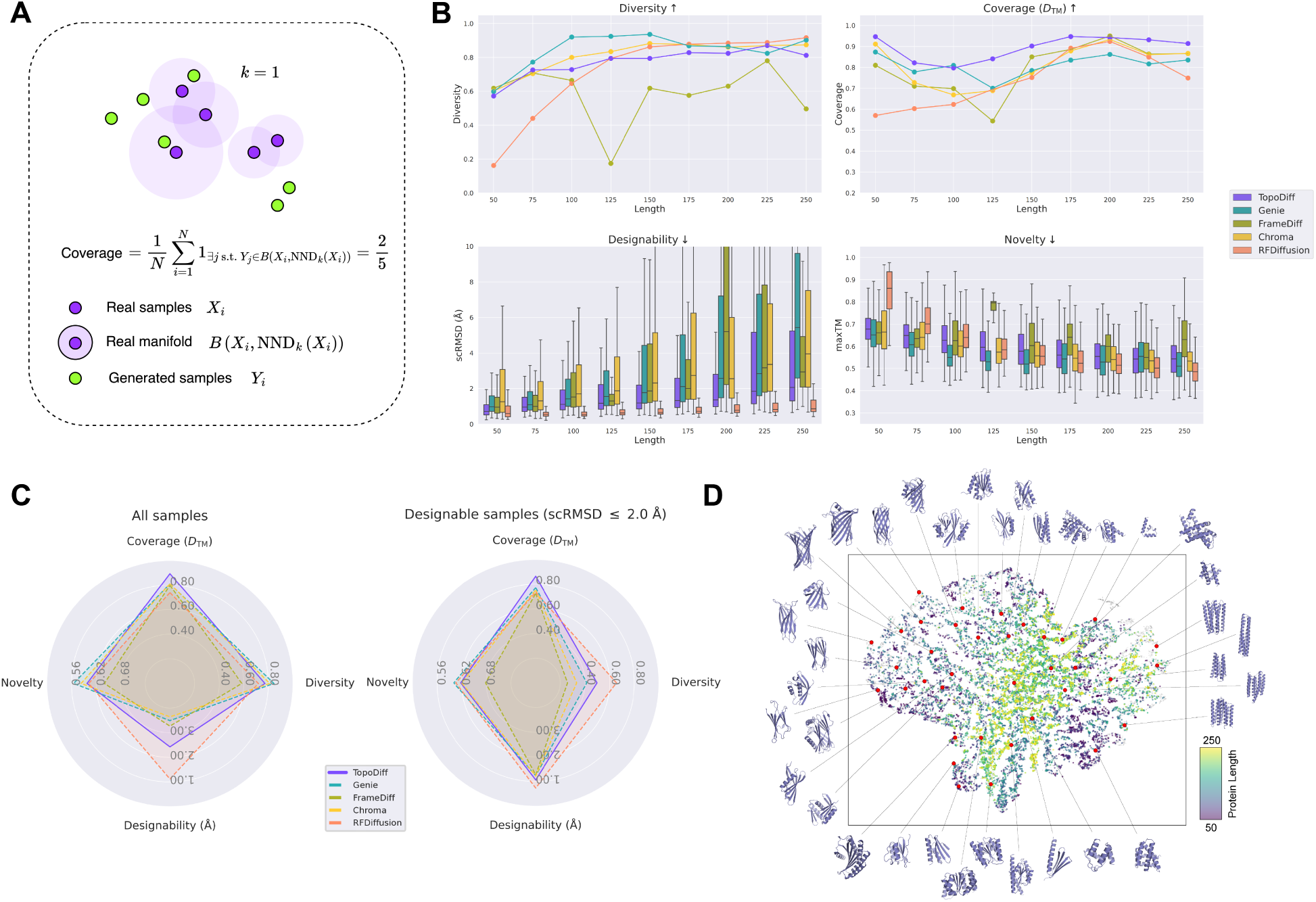
Evaluation of TopoDiff’s Generative Performance for Unconditional Sampling. **(A)** Illustration of the coverage metric used to quantify the extent to which the generated samples cover the natural protein fold space. Real samples *X_i_* are compared to generated samples *Y_j_* using a KNN approach to determine if each real sample is covered by generated samples within a defined distance threshold. The coverage score is calculated as the proportion of real samples that have at least one generated neighbor within this threshold. **(B)** Quantitative analysis of the generative performance across different models, measured by diversity, coverage, designability and novelty at varying protein lengths: 50, 75, 100, 125, 150, 175, 200, 225 and 250 residues. Arrows aside the metrics indicate the desired direction. **(C)** Radar plots summarizing the average performance of each model on different metrics (coverage, diversity, designability and novelty) when considering all generated samples (left) and only high-quality samples (right, scRMSD ≤ 2 Å). Each metric is averaged across all sampled lengths. **(D)** Projection of the sampled latent codes on the t-SNE dimension-reduced space of the CATH dataset. A total of 12,613 latent codes, corresponding to sampled structures with scRMSD ≤ 2 Å and maxTM < 0.7, are projected onto the original t-SNE space constructed from the CATH dataset (Figure 2). Each colored point represents a sampled latent code, with the color indicating the sampled length (50 ∼ 250, from purple to green). Representative structures generated from latent codes at different regions of the latent space are also visualized alongside the scatter plot, providing a visual inspection of the latent-structure relationship.

The first row of Figure 3B focuses on the evaluation of diversity and coverage. TopoDiff is comparable to the other methods in diversity, with the value gradually increasing from approximately 0.6 at the length of 50 to around 0.85 at lengths over 200. RFDiffusion shows a significant drawback in generating diversified structures at short lengths, despite its excellent power for longer proteins. In terms of coverage over the natural protein space, TopoDiff consistently outperforms all the other methods across the entire range of sampled lengths, demonstrating a robust performance. Along with its underperformance in diversity, RFDiffusion exhibits severely dropped coverage in the length range of [50, 150]. Similarly, FrameDiff also exhibits some degree of length-dependent instability on diversity and coverage. Genie excels at generating highly diversified samples, but still with a slightly lower coverage. To further investigate the specific advantage of TopoDiff over the others in the coverage of natural fold modes, we analyzed the sample-wise binary coverage indicators and found that our model could encompass a markedly larger number of natural folds with mainly beta-strand compositions, a group of topologies that are typically underrepresented by the other methods (Supplementary Figure S10).

The second row of Figure 3B shifts to the evaluation of designability and novelty. Regarding designability (in terms of scRMSD as defined in the “Methods” section), TopoDiff demonstrates advantages over most models excluding RFDiffusion, which has oversized parameters (Supplementary Table S1), across the entire sampled length range. Although RFDiffusion excels in generating credible designs, its reduced diversity and weakened coverage of the natural fold space in the short and medium length range of [50, 150] might limit its effectiveness in sampling novel folds, as more than 60% of natural domains in the CATH database settle in this range. As for novelty, we computed the maximal structural similarity (using TM-score^45^) of each individual sample to all structures in the CATH-40 database^20^ and plotted the distribution of all generated samples against the chain length. TopoDiff maintains a fair balance between the coverage of known folds and the generalizability to novel ones, with the median value of the maxTM (*i.e.* maximal TM-score) consistently around 0.6.

The model performance is further compared by averaging over all sampled lengths in Figure 3C, where the metrics are calculated using all samples (left subplot) and using samples with high designability (*i.e.* scRMSD ≤ 2 Å) only (right subplot), respectively. TopoDiff undoubtedly improves the overall coverage in both cases, indicating that the improvement is indeed attributable to truly designable samples. As mentioned in the “Methods” section, the computation of coverage relies on the definition of distance between a pair of structures (Equation 6). To supplement the coverage evaluation in Figure 3B&C where the average TM-score between query and target structures is uniformly used for the distance definition (Equation 7), we also present the result with the distances computed by a third-party model^46^ (Equation 8) in Supplementary Figure S9. Clearly, the overall relative trend among tested methods is similar for both distance definitions and the advantage of TopoDiff is consistent, indicating the robustness of our coverage computation upon the choice of distance definition. Overall, although TopoDiff samples with slightly lower designability than RFDiffusion, particularly at longer lengths, its designability still surpasses the other methods. Furthermore, TopoDiff generates samples at least three times faster than RFDiffusion (Supplementary Table S1), which enables the generation of more diverse and designable backbones within the same time period.

Finally, we seek to further explore the sampling space of TopoDiff. Out of the overall 22,500 latent codes and the resultant structures produced in this benchmark, we first filtered to retain latent-structure pairs with considerable designability and novelty (scRMSD ≤ 2 Å and maxTM < 0.7), resulting in 12,613 sampled pairs in total. We then projected the remaining latent codes onto the t-SNE subspace encoded by the CATH training samples (Figure 3D). To understand the relationship between the latent codes and the generated structures, we subsampled a group of latent codes spanning the entire latent space and co-visualized their corresponding structures around the scatter plot. As expected, the sampled latent codes show good coverage on the t-SNE dimension-reduced latent space, providing a strong foundation for the sufficient coverage of natural fold space by the sampled structures. Detailed inspection of the projected latent codes reveals that the latent samples are rather uniformly distributed on the manifold when the sampled length is below 150 but begins to favor the alpha-beta architectures such as the two-layer and three-layer sandwiches when the length grows, an observation that is consistent with the natural distribution of protein folds. A closer examination of the sampled structures shows that the spatial arrangement of secondary structure elements (SSEs) is highly correlated with the position of the conditioning latent code. On the rightmost side, the sampled structures are dominated by alpha bundles with several parallel helices. As the latent gradually shifts left, the helices begin to swing in different directions, forming orthogonal bundles. Then in the middle region of the manifold, beta sheets start to form adjacent to alpha helices, resulting in mixed sandwiches of different layers, alpha-beta barrels and/or more complex architectures. On the left side of the t-SNE manifold, beta sheets emerge as the predominant feature, forming rolls, barrels and beta sandwiches. Notably, this regional distribution pattern of structures aligns remarkably well with the original distribution of the annotated training samples (Figure 2B), which demonstrates that through the concurrent training scheme, not only does the encoder absorbs essential information of the global geometry into the latent encoding, but the diffusion decoder learns to utilize this information to generate structures with similar spatial characteristics.

In this section, we evaluate the fundamental capability of our trained latent sampler and structure decoder in unconditional structure sampling. In addition to the established metrics that emphasize the per-sample quality and within-sample diversity, we introduce a new coverage metric to quantify the diversity of fold modes in the sampled structures relative to natural structures, which helps to address the previously identified mode collapse issue^16^. TopoDiff, with its two-stage ancestral sampling design, achieves the best coverage, while delivering comparable or superior performance on the other metrics compared to models with similar parameter sizes. Visualization of the latent-structure pairs on the latent manifold demonstrates the unbiased sampling behavior of our model over the protein fold space.

### Controllable generation with the learned latent space

Beyond the common setting of unconditional sampling, the decomposition of the sampling process aforementioned in this study also introduces an entirely new dimension for imposing the condition and controlling the generation. Since the global geometry of sampled structures is primarily determined by the latent codes in our model, we can effectively control the conditional distribution of the structure space by manipulating the latent codes fed to the decoder (Equation 1). Controlled generation through the latent-level manipulation offers several unique characteristics that distinguish it from previous approaches. Unlike the discrete class labels that need predefinition to guide structure generation, the latent encoding used by TopoDiff requires no prior knowledge of the dataset partitioning and the continuity of the latent space further simplifies sampling. Additionally, the low-dimensional latent code provides a coarse-level constraint to the global geometry of a structure, which can be further refined by incorporating more detailed conditions to collectively control the generation process.

In Figure 4A, we illustrate how the different modules developed in this work can be integrated to enhance the controllability of structure sampling. The latent code could be sampled from the latent sampler shown in the previous section, or alternatively, harvested directly from the encoder’s posterior based on semantics. Moreover, additional latent classifiers can be developed to predict any properties desirable in the practical design and then used to tune the sampled latent distribution through classifier guidance or rejection sampling. Once a latent is determined, the structure sampling can be conditioned on both the latent code and additional residue-level information, achieving comprehensive constraining on both the global geometry and localized atomic details.

**Figure 4.**
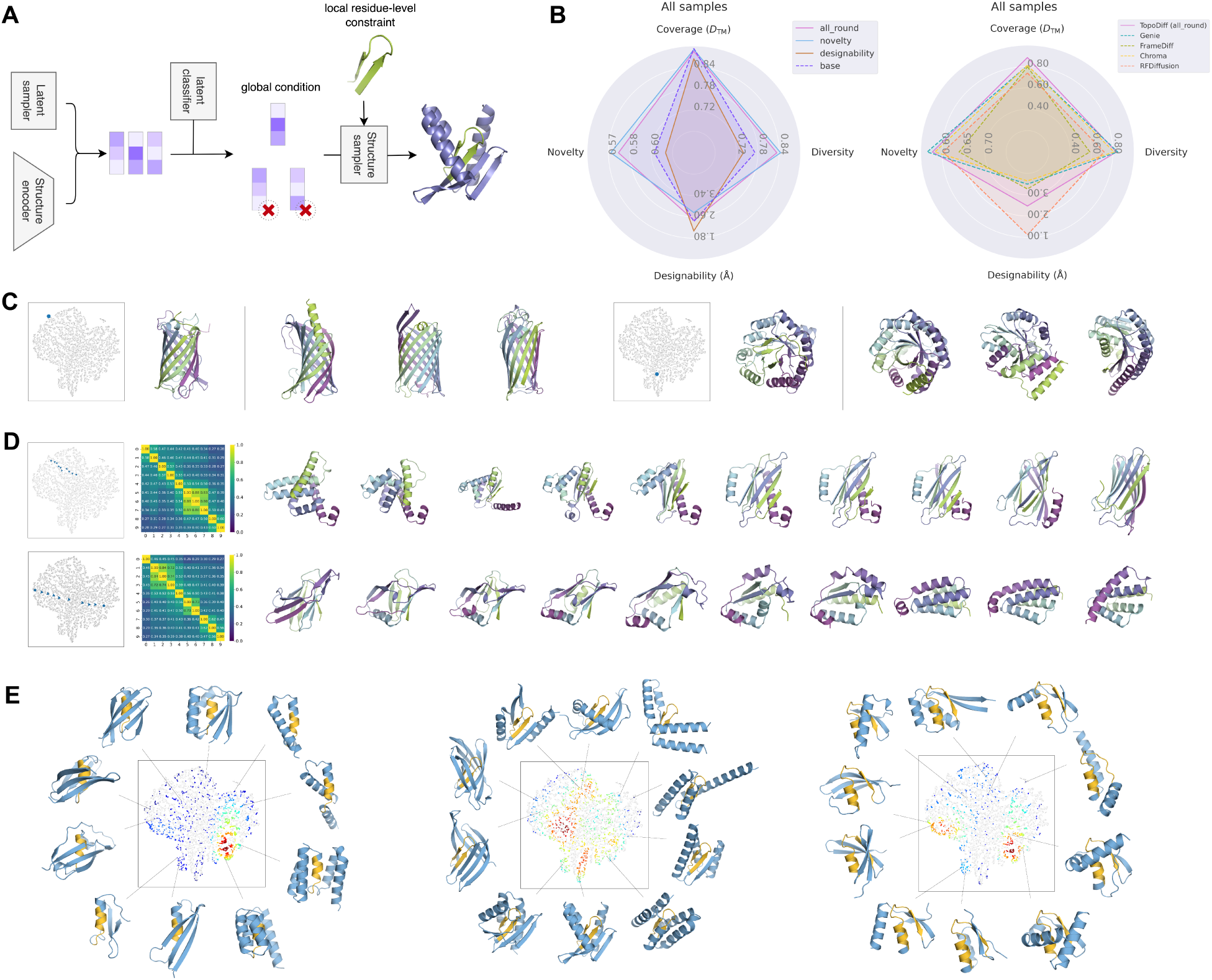
Exploring Controllable Protein Structure Generation with TopoDiff. **(A)** Overview of the controllable generation process. The latent code could be generated using the latent diffusion module or sampled from a local distribution. Downstream structure sampling is controlled by the acceptance ratio of the sampled latent codes. The accepted latent code, acting as a global condition, can be combined with local residue-level constraints to generate structures that meet specific design criteria. **(B)** Radar plots comparing the performance of different model variants derived from latent space rejection sampling: base, novelty-prioritized, designability-prioritized and all-round. In the left panel, the three model variants are compared with the base model. In the right panel, the all-round version of model variant is compared to other reference methods. All metrics are computed in the same way as Figure 3C. **(C)** Examples of structures generated by resampling based on the latent code of a reference structure. The leftmost structure represents the reference structure selected from the CATH dataset, with the right showing randomly generated structures without cherry-picking. **(D)** Visualization of structures generated by linear interpolation between two latent codes in the latent space. Each row depicts a different interpolation process, with the projected latent trajectory on the latent space (left), a TM-score distance matrix showing pairwise similarity between sampled structures (middle) and the visualization of these sampled structures (right). **(E)** Motif scaffolding experiment. For each motif case, latent codes corresponding to successful designs are plotted on the central scatter plot, with each point colored based on the kernel density estimation of successful designs in the local vicinity. Representative structures sampled from different regions of the latent space are shown around the scatter plot, with the query residue-level motifs highlighted in yellow and the rest colored blue.

Indeed, many measurable properties of the generated samples, like the proportion of SSEs, novelty and designability, exhibit specific spatial pattern in the latent space, due to their correlation with the protein global geometry in principle. For instance, some complex, mainly beta proteins are inherently more challenging design targets due to the stringent requirements for maintaining a global hydrogen bond network and avoiding improper tension under packing, which consequently leads to higher novelty but lower success rate, while at the other extreme, easier design targets like the up-down helix bundles often present extremely robust designability with weaker novelty. Hence, we explored the possibility of tuning the model performance by reweighting the sampling regions in the latent space. Specifically, we first generated sufficient samples across the entire latent manifold in the chain length of [50, 250], measured their scRMSD and maxTM values as defined in the benchmark, and then used these data to train simple multiple-layer-perception (MLP) prediction networks that forecast the expected desirable properties (*e.g.*, designability, novelty, SSE contents, etc.) for a given latent code. In the subsequent rounds of sampling, we readjusted the sampling probability of a sampled latent code based on its prediction results from these latent classifiers. This scheme effectively realizes the finetuning of model performance by simple manipulation at the low-dimensional latent level, which, unlike the exhaustive sampling at the atomic level in the Cartesian space, consumes negligible additional time.

Here, we attempted to tune the trade-off between designability and novelty through rejection sampling, where the sampled latent is accepted with a probability of 1 if it passes the latent classifier but with a probability of 0.1 otherwise (Equation 9). This strategy biases sampling towards the direction of enhancement in the desired property but still permits low-probability sample drawing from unfavored regions. We prepared three model variants, namely the designability-prior one, the novelty-prior one and the all-round one, which engage a designability classifier, a novelty classifier and a combination of both classifiers for rejection sampling, respectively. As expected, the designability-prior variant achieves elevated designability at the cost of decreased novelty, while the novelty-prior variant shows the opposite tendency (Figure 4B and Supplementary Figure S11). Interestingly, the all-round variant exhibits a balanced improvement over the base model, with novelty and diversity significantly increased while designability and coverage sustained.

In addition to general unconditional sampling in the latent space, the latent codes could also be sampled from any local region of interest. We first demonstrate the structure generation around a given query structure, based on latent sampling from a local distribution around its latent encoding. Since our latent encoder captures a few key degrees of freedom that describe the protein global geometry, repetitive sampling around the latent encoding of the query structure is supposed to produce an array of artificial structures sharing similar SSE spatial arrangements but with a greater variation in detailed residue connectivity. To test this capacity, we selected representative query proteins with a wide variety of architectures from the training dataset and randomly sampled 5 structures without further cherry-picking for each of them. Based on the the side-by-side comparison in Figure 4C and Supplementary Figure S12, the generated structures in general share a very similar SSE spatial arrangement to the query one with occasionally improved proportions of regular secondary structures, but always present a considerable level of diversity in the exact connectivity and topology. For architectures of higher levels of design difficulty, *e.g.*, the complex beta barrels, the original SSE architecture may deviate in some of the generated structures but is preserved in the rest.

Subsequently, we present the controlled structure generation based on interpolation between a pair of query latent codes. Indeed, the fixed-dimensional, continuous latent space learned by our model allows us to transit from one latent code to another through simple linear interpolation while ensuring all the inserted latent codes semantically meaningful. In this experiment, we collected a variety of latent pairs distantly located on the latent manifold and linearly interpolated 10 sequential intermediate latent codes for each pair. During structure generation, we adopted a slightly modified sampling strategy to ensure identical repeated sampling when conditioned on the same latent (see the “Methods” section). Figure 4D shows the structure variations along different latent trajectories, with each row representing the interpolation result for a distinctive latent pair. The second row presents the gradual change of generated structures from a mainly-beta roll to an all-alpha helix bundle, where each pair of adjacent structures in the trajectory share a certain degree of similarity (TM-score > 0.45), indicating an overall smooth transition process, although the two terminal structures have completely different SSE spatial arrangements and extremely low structural similarity (TM-score = 0.27). Notably, the adjacent structures attain a lower TM-score when the trajectory crosses the boundary of CATH classes in the latent space, and the beta strands always co-exist as hydrogen-bonded pairs in the generated structures, both observations consistent with intuitive perception and prior biophysical knowledge. The first row illustrates the interpolation from a structure of orthogonal helices to a beta sandwich, with particular attention paid to how the beta strand forms between the two helices and how the number of sheets gradually increases to form a roll and ultimately a sandwich. More cases could be found in Supplementary Figure S13.

Moreover, since the latent encoding provides a coarse control over the global geometry, it is possible to incorporate additional residue-level conditions to impose control at a finer level of granularity. We showcase this capacity through a motif scaffolding experiment. Specifically, we selected three representative motifs from the RFDiffusion paper^17^, including a single helix, a beta hairpin and a mixed pair of helix and strand. In this experiment, we first unconditionally sampled a series of latent codes that fit for the designed length, and then used both the latent and the specified motif as conditions to collectively generate the final structures. Figure 4E presents the density plots of all successful designs (*i.e.* scRMSD ≤ 2 Å, motif scRMSD ≤ 1 Å) on the latent manifold, along with representative sampled structures from different regions. In each case, the successful designs exhibit a non-uniform distribution across the latent manifold, achieving high populations in regions where the latent information and the motif are coherent. For instance, the single-helix 1YCR motif (left subplot) mainly succeeds at the manifold region dominated by orthogonal helices, while the two-strand 4ZYP motif (middle subplot) tends to generate designable structures in the alpha-beta sandwich region. When the structures inferred by the semantic latent conflict with the motif itself, *e.g.*, a helix in a predominantly beta region, the model seeks to find a compromised solution, albeit with increased difficulty. Taking the two-strand 4ZYP motif as an example, in the left part of the t-SNE-reduced manifold, we naturally sample many mainly-beta sandwich or roll-like structures. As the latent shifts right, alpha-beta rolls and sandwiches become more prevalent. Interestingly, even in the mainly-alpha region of the manifold, we occasionally sample successful designs where the two strands are either encompassed by a few long helices forming a stretched global shape (as shown in the top-right) or surrounded by several small helices forming a compact bundle (as shown in the bottom-right). Nevertheless, the latent code, when integrated with local constraints, acts as a global prompt to guide the model in exploring various architectures and topologies rather than always sampling from the preferred regions.

In this section, we discuss the unique advantages of our method in introducing new controllable structure generation schemes. We demonstrate that, with a simple and intuitive rejection sampling strategy, we can easily transform the model into different variants, one of which achieves equal or superior performance in all aspects compared to the base model. We then showcase controlled generation through two local latent sampling schemes, sampling around a query structure and interpolating between a given latent pair. We finally present the structure generation scenario of collectively controlling the global architecture and finer design objectives by combining the latent representation with localized detailed conditions. Noticeably, many of the these control mechanisms are feasible only in the model design proposed in this study.

### Generation of novel mainly-beta proteins and its experimental validation

In this section, we seek to test our TopoDiff model in the real-world design scenarios. We focus on the discovery of novel mainly-beta proteins, a large class of proteins that are commonly found in nature but remain underrepresented in *de novo* designed proteins. Due to the difficulties in precisely engineering non-local interactions and necessary structural irregularities (*e.g.*, beta bulges in beta strands) and in minimizing the aggregation propensity of exposed beta-strand edges, the *de novo* design of mainly-beta proteins lags far behind mainly-alpha and alpha-beta proteins^47^. Only a few experimentally validated cases have emerged in recent years^48–52^, which predominantly relied on predefined blueprints that were either extracted from naturally occurring proteins or based on parametric designing equations. While diffusion-based models have shown impressive potential in expanding the *de novo* design space of proteins, they have surprisingly not yet demonstrated superiority in generating experimentally validated novel mainly-beta proteins^17,22^. Given TopoDiff’s unprecedented coverage of the protein fold space, we are particularly interested in exploring whether it can generate novel mainly-beta proteins without the explicit use of human prior knowledge in designing detailed blueprints, and potentially push the boundaries of our understanding of the design constraints in *de novo* protein engineering.

We began by generating novel protein backbones through four rounds of sampling. For each round of sampling, we first sampled 22,500 backbones ranging from 50 to 200 residues, and applied a stringent filtering criterion to identify some of the most promising mainly-beta designs in terms of novelty and designability. In the first round of sampling, we conducted a completely unconditional sampling, resulting in 33 backbones with 70 sequence designs. In the subsequent three rounds, we gradually incorporated latent classifiers to reweight the sampled latent distribution based on the predicted beta ratio, novelty and designability. This approach effectively enforced *in silico* enrichment toward latent codes that were likely to decode into novel, designable mainly-beta proteins, yielding a fourfold increase in successful designs when using all three classifiers.Out of the final 403 backbones with 950 sequence designs, we further selected three designs per sampled length, resulting in 21 candidates for subsequent experimental validation. In Supplementary Figure S14, we visualize these candidates along with their novelty and designability metrics. These designs are highly diverse in their arrangement of beta strands, and exhibit notable novelty compared to known structures in PDB. All designs feature more than 50% of residues in beta strands and less than 20% in alpha helices, with more than half of designs consisting exclusively of beta strands and coils. Moreover, the global packing of these designs predominantly relies on the formation of beta sheets with numerous non-local interactions, making manual blueprint design exceptionally challenging. Regarding designability, all designs are predicted to be sufficiently foldable, as assessed by ESMFold and AlphaFold2. Notably, 16 out of 21 designs have all five AlphaFold2 models achieving pLDDT > 85% and scRMSD < 1.75 Å, which are considerably strong indicators of successful design^53^.

Following the *in silico* selection, we obtained synthetic genes encoding the 21 selected designs for subsequent wet lab experimental validation. Nine of these designs were expressed and soluble in *E. coli* (Supplementary Figure S15) and could be efficiently purified using nickel-affinity chromatography and size-exclusion chromatography (SEC) (Supplementary Figures S16A). SEC profiles of designs B07 and B10 exhibit distinct monomer peaks, while the others form a mixture of soluble aggregates and monomers (Figure 5A and Supplementary Figure S16B). Six of the nine expressed designs display the anticipated CD spectra for beta-sheet-rich proteins (Figure 5A and Supplementary Figure S16C). A summary of these experimental results could be found in Supplementary Table S2. In Figure 5A, we highlight four designs that attain the separable monomeric states and correct CD spectra. B07 and B10 demonstrate high thermal stability up to 95*^◦^*C, whereas B08 and B21 exhibit melting temperatures at approximately 80*^◦^*C and 65*^◦^*C, respectively. We further determined the structure of B10 with X-ray crystallography (Supplementary Figure S17 and Supplementary Table S3). As shown in Figure 5B, the solved structure closely matches the original backbone generated by TopoDiff, with a C*_α_*-RMSD of 1.31 Å.

**Figure 5.**
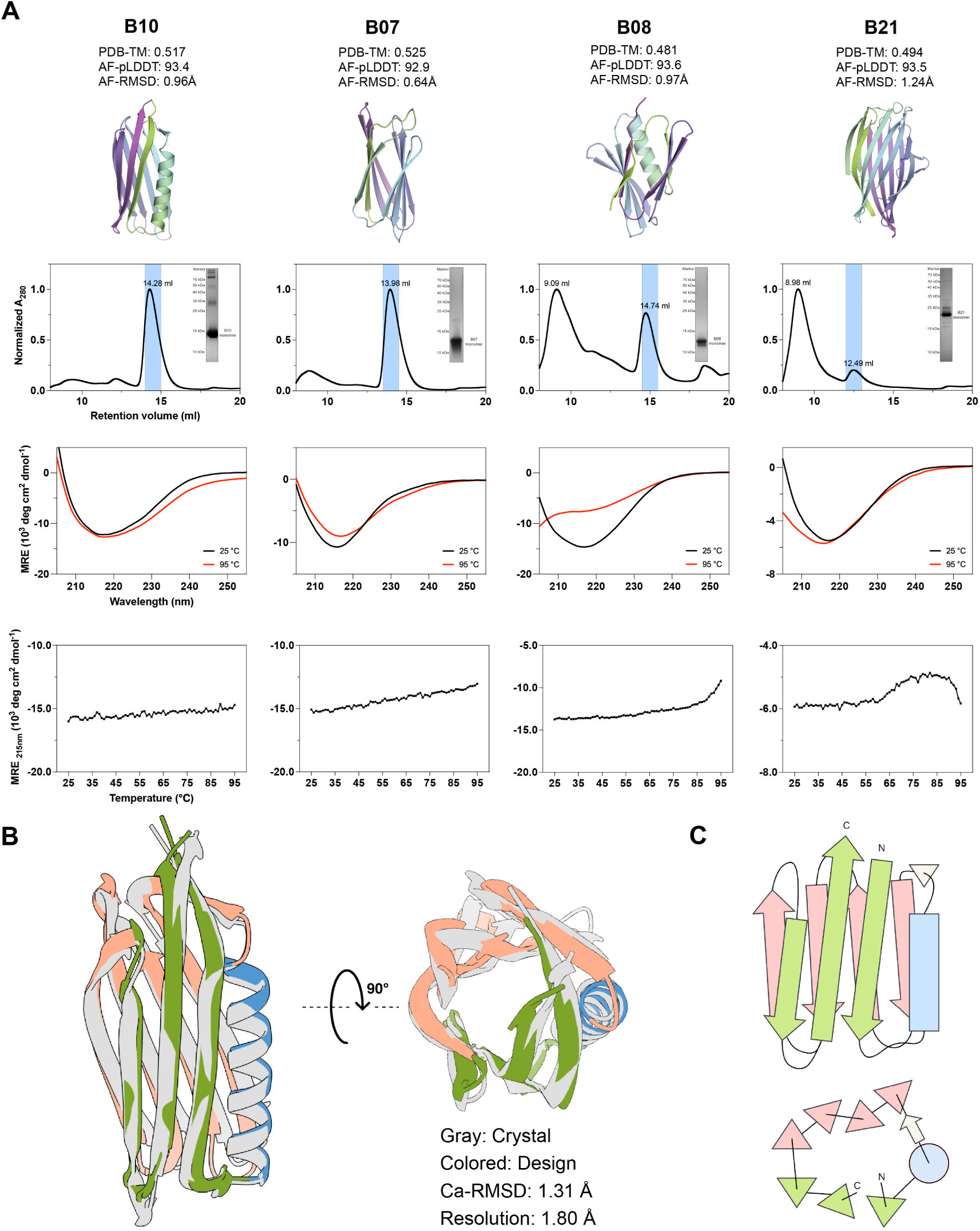
Experimental Validation of Novel Mainly-Beta Protein Designs. **(A)** Experimental characterizations of selected candidates (B10, B07, B08 and B21). The first row displays the design models with their corresponding metrics: PDB-TM (maximal TM-score to PDB), AF-RMSD (scRMSD compared to the best model of AlphaFold2) and AF-pLDDT (pLDDT of the best model of AlphaFold2). The second row shows the SEC profiles (with expected monomeric peaks highlighted in blue) and the monomer band observed on the SDS-PAGE gel. The third row presents CD spectra of the purified proteins at 25*^◦^*C (black) and 95*^◦^*C (red), demonstrating secondary structure content and thermostability. The fourth row displays the temperature dependence of the CD signal at 215 nm. No unfolding transition is observed at temperatures up to 95*^◦^*C for B10 and B07, suggesting their high thermostability. **(B)** The X-ray crystal structure (gray) of the design B10 matches closely with the generated backbone structure (colored) by TopoDiff (RMSD: 1.31 Å). **(C)** Topological diagrams of the design B10 in the side and top views.

All of the four successful designs shown in Figure 5A have the PDB-TM (*i.e.* maximum TM-score to PDB structures) lower than or close to 0.5, indicating their novelty in the SSE spatial arrangement and topology. Based on the solved X-ray structure, we focus on the design B10, a 125-residue mainly-beta protein composed of 8 beta strands and 1 alpha helix. One novel structural feature of this protein is the packing of the alpha helix into the crossover region between two beta sheets (Figure 5B), where the helix extends outward and pushes the adjacent beta strands of the two sheets apart, creating a unique triangular geometric arrangement in the top view (Figure 5C). Instead of a free-standing element, the alpha helix forms hydrophobic interactions with both beta sheets, contributing to the overall structural integrity. Interestingly, this compact triangular arrangement is unseen in natural proteins, with the closest structures in PDB being either beta barrels or two-layer sandwiches (Supplementary Figure S14). Additional analysis of the other monomeric designs is provided in Supplementary Results 7.

In summary, the above experimental results validate that our model, with its enhanced coverage of the fold space, is capable of generating new types of proteins that other models may struggle to sample, effectively generalizing beyond naturally occurring proteins to offer greater flexibility in designing novel structures tailored to user-defined objectives.

## Disccusion

In this work, we propose a novel framework that builds upon the current state-of-the-art diffusion generative model, enabling the concurrent learning of an encoder to capture a compact global structural representation and a conditional diffusion module to leverage this information for controllable generation. This improvement is achieved through a simple yet effective reformulation of the generation process into a two-stage hierarchical sampling method, the integration of a structure encoder and the additional modification for conditional constraining. Particularly, our approach does not rely on the intricate design of model architecture or the specifics of the diffusion formulation, nor does it require additional established labels or annotations, making it a highly versatile and easily implementable framework.

Unsupervisedly learning of the global structural representation in this study bypasses the dependence on existing classification systems and offers a novel continuous perspective of the fold space, which aligns well with established annotations across databases and provides additional levels of granularity. Introduction of this fixed-size latent encoding into the formulation of diffusion-based protein structure generation not only facilitates the human interpretation/understanding of the overall process, but also improves the coverage of the protein fold space while keeping the other performance metrics competitive. Notably, the latent space is intentionally confined to a low dimensionality with strong continuity, acting as a coarse constraining over the global geometry without hindering the discovery of novel folds during the generation process. This trade-off between the expressiveness of the latent representation and the generalizability to novel topologies is tunable for practical design objectives.

More importantly, the model design introduced in this study offers a new dimension for imposing conditions in the diffusion-based structure generation framework. We propose a few brand-new, versatile control schemes for protein structure generation through simple manipulation at the latent level. Furthermore, the latent-level control provides a useful supplement to the established residue-level conditions like the residue-wise SSE and pairwise adjacency^13,17^, which, albeit valuable, require substantial domain expertise and supposedly limit the sampling space. Hence, our method may benefit many practical design applications, particularly the scenarios where both global-level control and residue-level variation are desirable^51,54^.

In consideration of the unprecedentedly large scale of unannotated protein structure data revealed in the AlphaFold era, we intentionally avoid the usage of annotations on the training data, focusing on a more generic training scenario. Consequently, despite the current use of single-domain data with length ≤ 256 residues in the training set for simplifying comparison with established annotations, the whole framework operates in an unsupervised manner and is therefore scalable to the entire protein structure data with ease. Alternatively, this approach is customizable for a user-defined category of proteins, learning a class-specific representation to capture the inner variance and thus allowing for the development of a specialized generative model.

Based on the above discussions, we hope that our work, particularly its high-level design, will serve as a crucial contribution to the protein design community, inspiring further exploration and advancement in this domain.

In addition, in order to address the mode collapse challenge identified in previous works, we propose a metric to quantitatively evaluate the coverage of sampled structures over the established protein fold space. This metric offers flexibility in the exact implementation of the distance measurement, and we utilize two different definitions to validate its robustness. We anticipate that the development and adoption of this metric will facilitate the systematic evaluation of generative models in the field of protein design.

Eventually, we apply our method to a widely-recognized challenging design task: the design of mainly-beta proteins with novel backbone topologies. Our approach allows diffusion-based generation of mainly-beta or even full-beta novel proteins that have been validated by firm experimental evidence, without relying on any human pre-design. With this breakthrough, we prospect that new territories of the design space are available for exploration, such as the design of novel mainly-beta binders resembling natural binder proteins in the overall shape and interacting SSEs^41^. In spite of the successful cases presented, we note that a number of designs show clear aggregation tendencies (Supplementary Table S2), consistent with past experiences^47,49,55,56^. While deep-learning-based structure prediction models have proven useful in evaluating the designability of candidates in their monomeric folded states^53,57^, they still fall short in considering the aggregation propensity and folding dynamics^58,59^. With the progress in our work, consideration on these factors is expected to address the inherent difficulties associated with mainly-beta proteins, potentially leading to further amelioration of the success rate for their design.

## Methods

### The overall model design

In alignment with previous researches^13–17^, we represent the protein as a list of rigid transformations (residue clouds) in the SE(3)*^N^* space. Briefly, for a sequence of length *l*, each residue is parameterized as the collection of the translation of its C*_α_* atom, denoted as **x***_i_* ∈ *R*^3^, and its orientation, uniquely defined by the coordinates of three backbone atoms (C, C*_α_*, N) and denoted as **r***_i_* ∈ SO(3). Collectively, we denote the whole sequence as *R* = {(**x***_i_,* **r***_i_*)} ∈ SE(3)*^l^*.

A distinctive step we took as compared to the previous works^13–17^ is the introduction of a fixed-size, low-dimensional latent variable **z** ∈ *R^C^***^z^**, which encodes the essential information about the global geometry of the underlying structure. We further model the joint distribution *p* (*R,* **z**) in a hierarchical fashion. With this formulation, we essentially decompose the generation process into two steps, namely the acquisition of **z** and the subsequent structure space generation process conditioned on it:

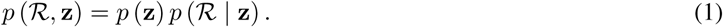

The conditional generation in the structure space is further modeled as a diffusion-based process, through which our model is trained to learn the complex data distribution. In brief, we define the forward process of the diffusion model as a non-learnable Markovian process that gradually introduces noises to a protein structure *R*^0^ towards a pre-defined prior distribution *p_T_* within *T* steps

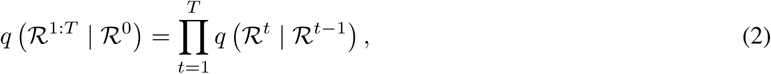

while the reverse process is also a Markovian process that learns to remove the noise signal conditioned on the latent variable **z**

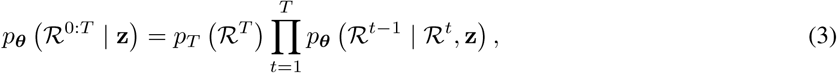

where *q* and *p* are the probability distributions for the forward and reverse processes, respectively, with *θ* denoting the model parameters. The training objective is to make an accurate prediction 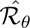 for the ground truth *R*0, which is, for simplicity, denoted here as the reconstruction loss 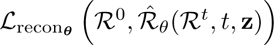.

Although the current protein structure classification systems^19,20,32,60^ partially align with our definition and expectation of **z**, several notable drawbacks preclude them as the optimal representations. First, all classification systems implicitly assume protein folds as discrete islands in the structure space, but overlook the connections between these discrete collections^39^. To enforce this discreteness in annotation assignments, the systems must rely on subjective and arbitrary criteria, which often lead to inconsistencies between systems and frequently cause confusion and controversy^24,36,39^. Second, the explicit requirement of manual annotations negates the possibility of scaling up to the whole known structure space or repurposing to some smaller and specialized dataset, where the annotation is lacking and the learning of representations is still an intriguing question.

To tackle this challenge, we seek to learn a continuous representation of **z** directly from the training data. Indeed, alternative to a discrete view, some past studies also suggested that the complete fold space, and consequently the space of *de novo* designed proteins, should be considered geometrically continuous^23,38,61^. In light of past attempts^62–66^, we incorporate an SE(3)-equivariant structure encoder (with parameters *ϕ*) to infer **z** from the input ground truth structure following the posterior distribution *q****_ϕ_***(**z** | *R*^0^). We model the *p*(**z**) with an isotropic Gaussian prior, and add a Kullback-Leibler (KL) divergence loss term to encourage a continuous latent structure, effectively shaping our model into a VAE-like framework. Combined with the diffusion model, our final training objective is

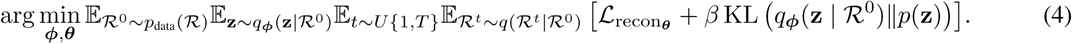

With this training scheme, we concurrently trained an encoder to capture the essential information about the global geometry of a structure and a decoder to sample in the structure space by conditioning on it. The considerably low dimensionality of the latent and the distribution regularization effect (by the KL divergence loss) jointly force the encoder to capture several crucial degrees of freedom that account for the variation within the dataset, simplifying the latent distribution. By encoding all training samples into the latent space, we further trained a latent diffusion model to capture this distribution and eventually achieve unconditional sampling of the structure space *p* (R) through the unconditional sampling of *p* (**z**) and the conditional generation of *p* (*R* | **z**) (Equation 1).

### Dataset and training summary

We prepared two datasets for the training of TopoDiff: the PDB monomer set and the CATH-60 set. PDB monomer set was directly collected from the PDB^18^, and the CATH-60 dataset was constructed based on the S60 non-redundant domain list from the CATH 4.3 release^20^. For the training of the structure diffusion module, we employed a strategy that considers both model performance and training efficiency. Specifically, we first trained the structure diffusion module alone on the PDB monomer set to learn a good generative prior to the protein structure space, and in the later phase concurrently trained this base model with a randomly initialized structure encoder in the aforementioned architecture.

### Visualization of structure representation of different databases

We collected domain-level classification datasets from three different sources: CATH^19,20^, SCOPe^32^ and ECOD^33^. The structure encoder was trained exclusively on the CATH dataset. The other two datasets, SCOPe and ECOD, were used solely during the inference stage, and we restricted the included structures to single-chain proteins with a maximum of 256 residues.

To get the representation of each structure, we extracted the coordinates of C*_α_* atoms and inferred through the trained encoder. For dimension reduction, we applied the t-SNE^30^ algorithm to compute the transformed 2D embeddings of the latent representations. Specifically, we used the implementation from openTSNE^67^, with the L2 distance metric and a perplexity of 50.

To visually assess the agreement of the learned latent space with human annotations, we colored the t-SNE scatter plot according to different annotation hierarchies. For the top hierarchy of each database, we used a discrete colormap to differentiate categories with distinct colors. For the second hierarchy, such as the Architecture level in the CATH database, we created subplots for each architecture category, coloring the samples according to the kernel density estimation^68,69^ of that category on the 2D latent space, providing a clear visualization of each category’s distribution.

### Computation and visualization of empirical latent distribution distance between CATH architectures

To get the difference between the empirical latent distributions of two CATH architectures, we first grouped the pre-computed latent representations according to their respective architectures and then calculated their Wasserstein distance in the latent space. Given two sets of points in the latent space, 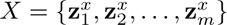 and 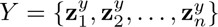, the Wasserstein distance, also referred to as the Earth Mover Distance and abbreviated as EMD here, is defined as:

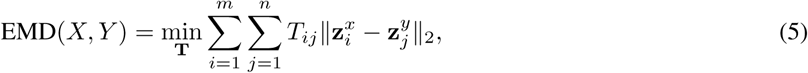

subject to the constraints

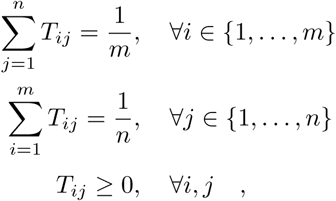

where *T_ij_*s are the parameters awaiting optimization and ∥ · ∥_2_ refers to the L2 norm.

We used the Python Optimal Transport (POT) library^70^ to compute the EMD with the ot.emd2 function. After calculating the pairwise distribution distances between all architecture pairs, we constructed a graph diagram to illustrate the continuous relationships among the major groups. We included only architectures with more than 200 structures and preserved edges with EMD < 1.5. To ensure that the graph diagram accurately reflects the distance relationships while maintaining a clear visual clarity, we used nx.spring_layout to optimize the layout with k = 1 and iterations = 100.

### Identification of latent outliers from different structure categories

Based on the latent encoding, we could identify outliers in the structural categories, *i.e.* proteins that are either misclassified or ambiguously classified, the latter of which refers to proteins that are likely to exhibit structural features of more than one categories simultaneously. To identify such proteins, we calculated the averaged Euclidean distance between each protein and all proteins within the same category in the latent space, and ranked proteins by their mean intra-category distances to identify the outliers within each category. For the aforementioned identification, we conducted an analysis on the first two hierarchies of the CATH dataset, namely the class and architecture levels, among which the former is particularly notable because CATH has only three classifications at this level. To ensure that the identified proteins are indeed misclassified or ambiguously classified rather than merely structurally unique proteins, we added the information of neighboring proteins in the latent space as an additional selection criterion, requiring that the target has more than 60% of its 20 latent-space nearest neighbors belonging to a category different from the CATH assignment.

### Definition of evaluation metrics for structure generation

To comprehensively benchmark the quality of structures generated by various diffusion models, we evaluated the samples with a series of metrics, each emphasizing a distinct aspect.

#### Coverage

we adapt the coverage metric initially proposed by Naeem et al.^44^ to measure the extent to which a model can cover the natural protein space. Briefly, we first constructed a KNN manifold of all real samples (*i.e.* natural proteins) and then measured the fraction of real samples whose neighborhoods encompass at least one fake sample (*i.e.* structures produced by the generative models). Formally, for *N* given samples, the coverage over the target sample distribution is defined as:

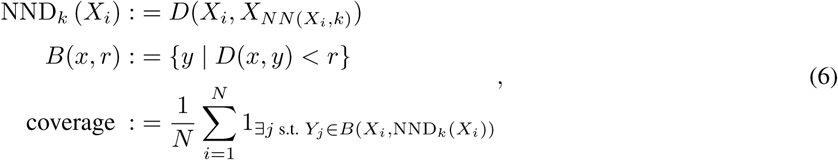

where {*X_i_*} denotes the set of real samples, {*Y_i_*} denotes the set of all fake samples, NND*_k_*(*X_i_*) stands for the distance from *X_i_*to its *k^th^*nearest neighbor in {*X_i_*} excluding itself, and *B*(*x, r*) refers to the hypersphere around *x* with a radius of *r*.

Based on this definition, we need to define a function *D*(·, ·) to measure the distance between two arbitrary structures. Specifically, we need to compute at least the distance between *X_i_* and the *k^th^* nearest neighbors in {*X_j_* | *j*≠ *i*} to construct the KNN manifold at the given point, and then use the distance between *X_i_* and its 1*^st^* nearest neighbor in {*Y_i_*} to decide whether *X_i_* is covered by an arbitrary fake sample. The final coverage metric is obtained by averaging the binary indicators over {*X_i_*}. In this work, we computed the metric with two different distance definitions to demonstrate the flexibility and robustness.

The first definition is the complement of TM-score^45,71^ of two compared structures. To make the function symmetric with respect to the order of inputs, we define it based on the average of query TM-score and target TM-score:

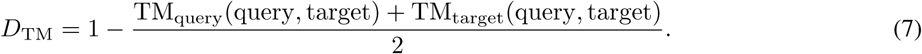

This distance definition is a natural choice as the TM-score is designed to measure the global structural similarity and has been widely used for structure evaluation^72^. However, due to its use of dynamic programming and heuristic iterative algorithms to refine for optimal solutions, the computation of this metric is generally non-parallelizable and will be exceptionally slow when comparing a sampled structure to all natural structures. Therefore, we also provide a faster-speed alternative, taking the distance defined by a third-party structure searching model^73^, which employs supervised contrastive learning to learn an embedding vector (**emb**) of each protein for structure comparison:

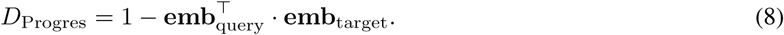

In practice, the pairwise distance between natural structures is pre-computed with both approaches for reuse.

The choice of the hyperparameter *k* of KNN should be determined prior to the computation of the metric. Following the recommendation of the original paper by Naeem et al.^44^, we chose *k* based the principle that a sample size equivalent to the artificial samples from the very same distribution of real samples could achieve a coverage close enough to 1. To do this, we initially randomly sampled 500 natural chains as the pseudo-query set, used the remaining chains as samples from the target distribution, and then computed the coverage of the pseudo-query set against the target distributions for different choices of *k*. Based on such a scheme, we ultimately used *k* = 100 for all experiments, although we found that different choices of *k* generally do not alter the relative rankings of the evaluated models (Supplementary Figure S8).

When comparing the samples of a fixed length to the natural protein distribution, at each sampling length *l*, we considered all natural protein structures in the CATH-40 dataset^20^ lying within the interval of [*l* − 25, *l* + 25].

#### Diversity

To compute the diversity of *N* samples, we first used TM-align^71^ to compute the pairwise TM-scores. Then we clustered the samples with a cutoff of 0.6. The proportion of total clusters to the total number of samples *N* was reported as diversity: a higher score generally indicates that the generated samples are more diverse.

#### Designability

To assess the designability of a given sample, we first used ProteinMPNN^74^ to sample 8 amino acid sequences with a temperature of 0.1. Subsequently, the sequences were fed to ESMFold^75^ to infer the structures. The minimum RMSD (*i.e.* root mean square distance) of the inferred structures to the given sample was reported as scRMSD (*i.e.* self-consistent RMSD): a smaller value generally implies that the sample is more designable.

#### Novelty

To assess the novelty of a given sample, we began by using Foldseek^76^ to query the sample against the CATH-40 dataset^20^ with the parameter “-a 1 –exhaustive-search 1 -e inf -c 0.5 –alignment-type 1”. As Foldseek employs a slightly different implementation of TM-align^71^, we subsequently selected the top 25 matches from the query results with the highest TM-scores and recomputed the alignment with TM-align^71^. The highest TM-score to the chains in the dataset was reported as maxTM (length-normalized by the sampled structures), representing the novelty of a sample: a higher score generally implies that the sample is less novel.

In the experimental validation section, to assess novelty against the full PDB scope, we used Foldseek^76^ to query the entire PDB (data up to February 2023) with the parameters “-a 1 –exhaustive-search 0 -e inf -c 0.5 –alignment-type 1” (with exhaustive pairwise search disabled). The top 25 matches were then realigned using TM-align, and the highest query TM-score was reported.

### Benchmark on unconditional sampling

All benchmark experiments were conducted with TopoDiff Model v1.1.2. For each model evaluated in the benchmark testing, we randomly sampled 500 structures at each length of {50, 75, 100, 125, 150, 175, 200, 225, 250}. We first computed metrics based on all 500 samples at each length, to get a series of values reflecting the length-dependent performance for each model. Since some sampled structures tended to exhibit considerably low designability and, in the meantime, high novelty and diversity due to the structural defects, leading to an overestimation of these metrics, we also implemented an additional step to filter the samples by only preserving those with scRMSD ≤ 2 Å and recomputed the metrics at each length. Finally, for each metric, we averaged out along all sampled lengths to present an indication of its overall performance. We used the length-averaged metrics to draw the radar plot as shown in Figure 3C.

### Co-visualization of latent space and generated structures

To project the latent codes of sampled structures onto the t-SNE dimension-reduced manifold, we used the transform method implemented in openTSNE^67^ to embed the newly sampled latents into the same space we used for CATH dataset visualization. Briefly, following the same basic principles as conventional t-SNE, the positions of the background embeddings were kept fixed and each sampled latent was optimized independently with respect to them. We only selected the latent codes associated with the fairly designable and novel samples (*i.e.* scRMSD ≤ 2 Å, CATH-maxTM < 0.7). Finally, we also co-visualized some sampled structures spanning the entire latent manifold, providing a visual context on the sampled structures and their spatial relationships with respect to the latent codes.

### Controlling model preference with the latent-space rejection sampling

To tune the sampled latent distribution, we began by training latent classifiers that predict the structural properties of given latent codes. We further used these latent classifiers to conduct rejection sampling over the sampled latents, obtaining variants of the model with distinct sampling preferences. Recall that *p_θ_*(**z**) is the base distribution we unconditionally sample from the latent diffusion module. Our goal is to reweight the sampled latent distribution using a set of trained classifier functions *f*_1_(**z**), …, *f_n_*(**z**) and corresponding thresholds *c*_1_*, …, c_n_* based on an acceptance probability function *α*(**z**), such that the resulting latent distribution *p*_accept_(**z**) (and eventually the sampled structure distribution) would be shifted towards our intended preference.

Specifically, for each variant, we first defined a series of length-dependent thresholds in terms of the designability (scRMSD) and novelty (maxTM) of the sampled structures. For simplicity, we only considered three possible variants: designability-prior (sampling highly designable structures with the designability classifier), novelty-prior (sampling highly novel structures with the novelty classifier) and all-round (sampling structures with fairly high designability and novelty when collectively using both classifiers). The exact thresholds we used at various sample lengths are summarized in Table 1.

**Table 1.** Thresholds used for different model variants during rejection sampling.

| Model variant | Designability threshold | Novelty threshold |
| --- | --- | --- |
| **designability-prior** | [0.3, 0.3, 0.35, 0.35, 0.35, 0.35, 0.35, 0.35, 0.35] | — |
| **novelty-prior** | — | [0.66, 0.66, 0.6, 0.6, 0.56, 0.56, 0.55, 0.55, 0.55] |
| **all-round** | [0.5, 0.5, 0.5, 0.5, 0.5, 0.6, 0.6, 0.6, 0.6] | [0.66, 0.66, 0.6, 0.6, 0.56, 0.56, 0.55, 0.55, 0.55] |

Intuitively, we want to sample the latent codes with a higher probability when they match the expectation, and lower if not. To achieve this aim, we define an acceptance probability function *α*(**z**), requiring that **z** is accepted with the probability of 1 only if approved by all classifiers and with the probability of 0.1 if denied by any classifier:

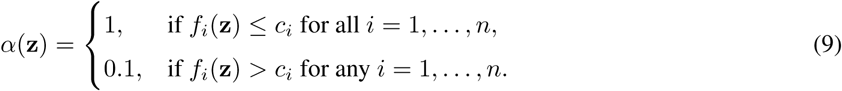

The resulting probability density function after applying the rejection sampling procedure is

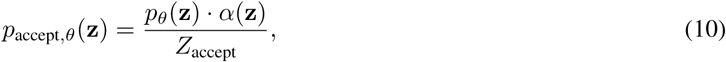

where:

– *p_θ_*(**z**) is the original latent distribution sampled from the latent diffusion module,
– *α*(**z**) is the combined acceptance probability as defined above,
– and *Z*_accept_ is the normalization constant, ensuring that *p*_accept_*_,θ_*(**z**) integrates to 1.

This formulation ensures that the resulting distribution *p*_accept_*_,θ_*(**z**) reflects the collective influence of all the classifiers. By training all modules once and setting different combinations of these thresholds, we shifted the sampled distribution to match our expectations as closely as possible.

For each variant, we sampled 500 structures per sample length using the given latent sampling strategy, and computed all metrics following the definition introduced in the benchmark section.

### Sampling of structures with similar global geometry

To sample structures sharing similar global geometry with respect to a reference structure, we encoded the query structures in our CATH training set using the trained encoder, and randomly resampled 5 structures with each inferred latent code. We selected reference structures with a wide variety of their SSE compositions and spatial arrangements, and listed their original structures and the five non-cherrypicked samples side by side.

### Sampling of structures with latent code interpolation

For the interpolation experiment, we first randomly chose a number of candidate pairs of samples from the CATH training set. For each candidate pair, we conducted a linear interpolation to get 10 latent codes evenly located between the two termini. Finally, we randomly selected a chain length between 75 and 150 residues, and sampled a structure with each latent code.

For structure sampling, we implemented with a setting that minimizes the stochasticity introduced into the reverse sampling process. Specifically, in the R^3^ space, we used DDIM (Denoising Diffusion Implicit Models)^77^ formulation instead of the default DDPM one, where the reverse sampling process is reformulated in a noise-free way that *R^t−^*^1^ is deterministically derived given *R^t^*, 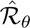 and the noise schedule. Similarly, in the SO(3) space, we used a score scale of 1 or 2 and a noise scale of 0, such that the reverse sampling process would also introduce no additional stochasticity. By using this reverse sampling setting, we ensured that the generated structures are solely determined by the initial state of noise sampled from the SE(3)*^l^* space and the conditioned latent code. For the structure generation of each series of interpolated latent codes, we further fixed the random seed used for the initial state sampling, so that the the only difference between the generation of each structure is the latent code itself. By this means, we could ensure that the generated structures are more consistent with the latent code they are conditioned on and reduce unwanted variability caused by sampling stochasticity.

### Sampling of structures with motif scaffolding

For the motif scaffolding experiment, we selected three representative motifs from the RFDiffusion work^17^, whose original structures adopt distinct secondary structures (mainly alpha, mixed of alpha and beta, and mainly beta). We largely followed the experiment setting introduced in the original publication^17^. Specifically, each design case was first associated with a detailed setting on the motif information and constraints, such as the total length of designs, the motif definition and the sequential position of motif on designs. Subsequently, we randomly sampled 5,000 valid combinations of constraints and latent codes, and used these combinations for structure sampling. For each generated structure, we designed 8 sequences with ProteinMPNN^74^ and folded them with ESMFold^75^. The designs were marked successful if the global scRMSD ≤ 2 Å and the motif scRMSD ≤ 1 Å. We kept these settings as close as possible with respect to the original publication^17^, except two differences. First, for the two-strand 4ZYP motif, since the originally designed length (30 ∼ 50) was too short to adopt most of the stable mainly-beta topologies, we kept the motif fixed and increased the sample length to 90 ∼ 110. Second, during sequence design, we did not freeze the motif sequence as our structure diffusion module was not specifically trained to be aware of the motif sequence information. The final motif design constraints are summarized in Table 2.

**Table 2.**
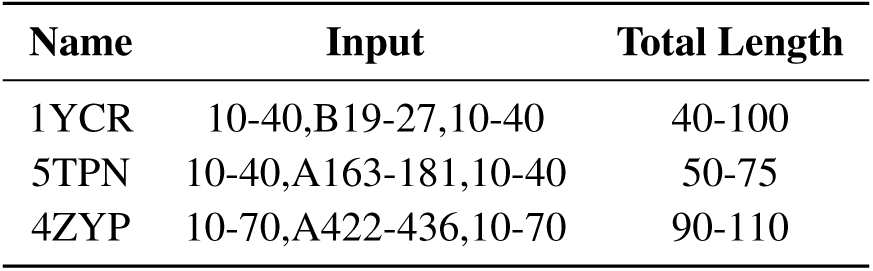
Information of motif design tasks.

| Name | Input | Total Length |
| --- | --- | --- |
| 1YCR | 10-40,B19-27,10-40 | 40-100 |
| 5TPN | 10-40,A163-181,10-40 | 50-75 |
| 4ZYP | 10-70,A422-436,10-70 | 90-110 |

For each motif, we gathered all successful designs and projected the corresponding latent codes onto the t-SNE dimension-reduced plot. The scatter plot is visualized as a heatmap, with each point colored according to the kernel density estimation^68,69^ of successful designs on the t-SNE space. We also co-visualized generated structures from each region of the manifold to demonstrate the diversity and controllability on the global geometry. In the structural presentation, the motif part is colored in yellow and the rest in sky blue.

### *In silico* design of novel mainly-beta proteins

Due to the asynchronous development of the model, all the experiment validations in this section were conducted using TopoDiff Model v1.1.1. All supplementary modules (*e.g.*, latent diffusion module, latent classifiers, etc.) used in this experiment were also trained based on this version. Briefly, we first sampled a number of protein backbones using the aforementioned rejection sampling strategy to focus the sampling on mainly-beta proteins with good novelty and designability. We then adopted a three-stage filtering pipeline to generate sequence designs and screen for designs with exceptionally good *in silico* quality. Finally, out of the 403 resulting backbone with 950 sequence designs, we selected 21 designs ranging from 50 residues to 200 residues for experimental validation.

### Protein expression and purification

Genes encoding the selected designs were synthesized from GenScript, cloned onto pET-29b(+) vector using GoldenGate assembly, verified with Sanger sequencing, and transformed into the BL21 *E. coli* strain. Protein expression was induced in the TBM-5052 medium with 50 *µ*g/mL kanamycin overnight at 37*^◦^*C. Expression was evaluated through Western blot against the N-terminal His tag. Cells with expressed designs were lysed by sonication for 0.2-L scale cultures and centrifuged at 12,000 g for an hour. The supernatant was purified using immobilized metal affinity chromatography with the Ni-NTA affinity resin.

### Size-exclusion chromatography (SEC)

Eluted protein samples were concentrated using AmiconUltra 3 kDa concentrators with a final volume of 1.2 mL, and then applied to a Superdex75 increase 10/300 GL column on an ÄKTA avant 25. Analysis of each component during the process was performed using sulfate-polyacrylamide gel electrophoresis (SDS-PAGE). Monomeric peaks were collected for further experiment or frozen for further experiments.

### Circular dichroism (CD)

Purified proteins were diluted at 0.2 mg/mL in the PBS buffer (25 mM phosphate, 150 mM NaCl, pH = 7.4). The concentrations were determined from the absorbance at 280 nm and the relative extinction coefficients. Wavelength scans from 190 nm to 280 nm were recorded at 25*^◦^*C and 95*^◦^*C using a 1-mm pathlength cuvette on a CD spectrophotometer. Background spectra were acquired across the same range and manually subtracted after conversion. Thermal denaturation was monitored at 215 nm from 25*^◦^*C to 95*^◦^*C. Mean residue ellipticity (MRE) [*θ*] was calculated from the original CD data (in mdeg) and was normalized by the sample concentration and the sequence length.

### Crystallization and structure determination

Crystallization screening was performed using commercially available kit sets through the sitting drop vapor diffusion at 16*^◦^*C. Diffraction-quality crystals were obtained for the design B10 with equal volume reservoir buffer (30% PEG-3350 (w/v), 0.1 M Tris-HCl, 0.2 M NaCl, pH = 8.5). Crystals were flash cooled in the liquid nitrogen without cryoprotection. Data were collected on 02U1 at the Shanghai Synchrotron Radiation Facility (SSRF). Data images were processed using the program HKL2000^78^. Crystal structures were solved by molecular replacement using the Phenix^79^ software suite with AlphaFold2^11^ prediction as the search model. The structure refinement was also performed using Phenix. The program COOT^80^ was used for manual rebuilding, while PyMOL^81^ and ChimeraX^82^ were used for generating the molecular graphics images.

## Data availability

The dataset used for model training, along with the trained model weights, benchmark data and protein designs selected for experimental validation, are available at https://zenodo.org/records/13879812.

## Code availability

The TopoDiff model is implemented in PyTorch. Full scripts and guidance for utilizing the model are available at https://github.com/meneshail/TopoDiff/tree/main. The complete training script will be made available upon acceptance of the manuscript.

## Author contribution

Y. Z. and H. G. conceived the study. Y. Z. and Z. M. designed and implemented the model. Y. Z. and Z. M. performed *in silico* experiments and analyzed the results. Y. Z. designed candidate proteins for experimental validation. Y. L. designed, executed and analyzed all wet-laboratory experiments. H. G. supervised the development of the model and result analysis. C. X. supervised the design of candidate proteins and wet-laboratory experiments. M. L. contributed to the X-ray structure determination. Y. Z. drafted the initial manuscript. Y. Z., Y. L. and Z. M. created the final figures. All authors contributed to writing and improving the manuscript, and approved the submission.

## Acknowledgments

This work has been supported by the Ministry of Science and Technology of China (#2023YFF1204400), by the National Natural Science Foundation of China (#32171243), and by the Beijing Frontier Research Center for Biological Structure. We thank the staff of Beamline BL02U1, BL10U2, BL18U1 and BL19U1 at the Shanghai Synchrotron Radiation Facility as well as X-ray crystallography platform, National Protein Science Facility, Tsinghua University, for the assistance in X-ray diffraction data collection and analysis. We thank Jian Hu, Zefeng Zhu, Yi Xue and Chen Song for helpful discussions.

## Supplementary Information

### Supplementary Results

#### 1 Additional results for latent space visualization

**Figure S1.**
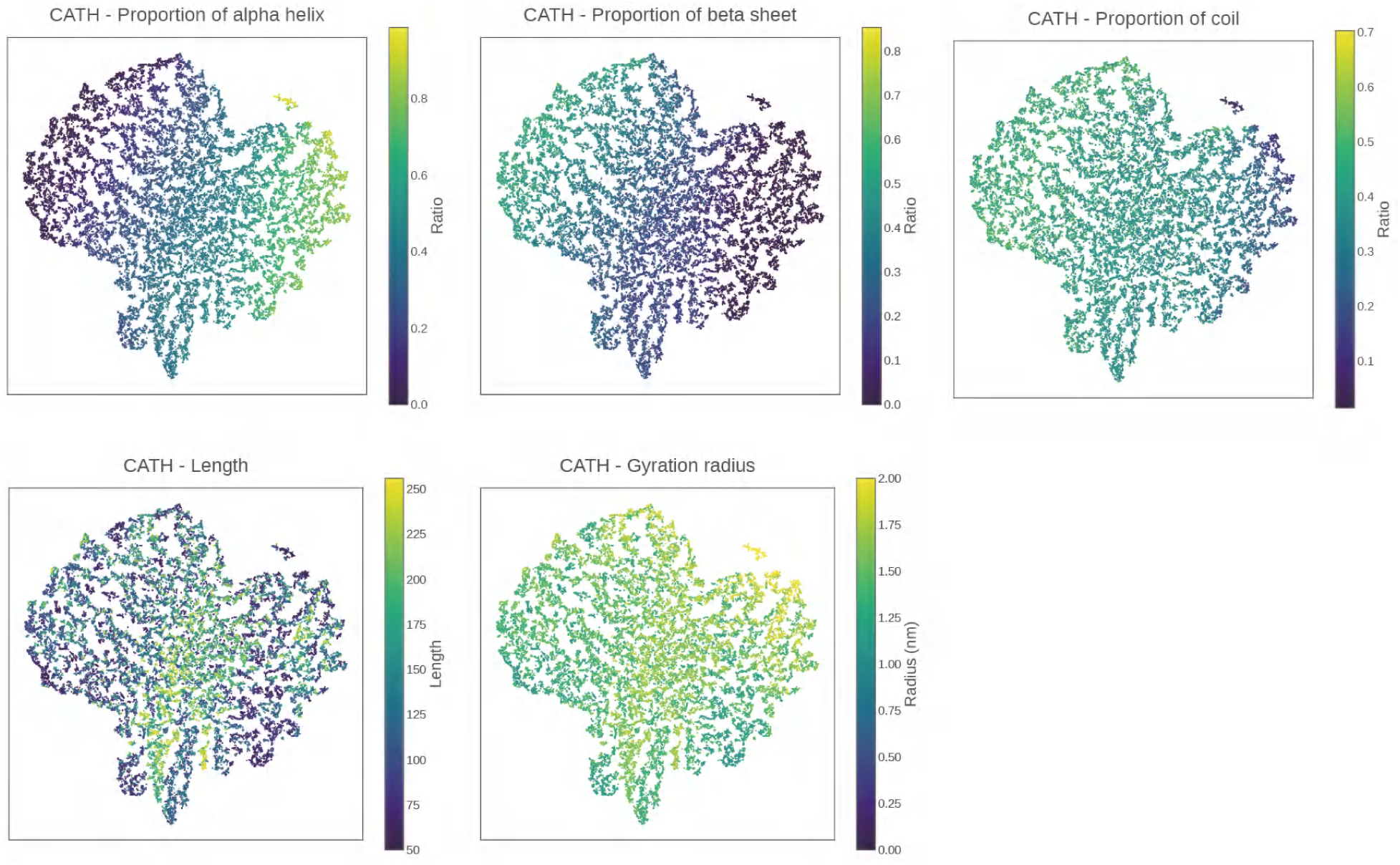
Visualization of inherent structural descriptors on the latent space. The plots display various structural descriptors mapped onto the latent space for all protein samples in the CATH dataset. Commonly used descriptors, such as the proportions of the alpha-helix, beta-sheet and coil secondary structures, along with the chain length and the radius of gyration, are visualized. In each subplot, the protein samples are colored according to their respective values of the corresponding descriptor.

**Figure S2.**
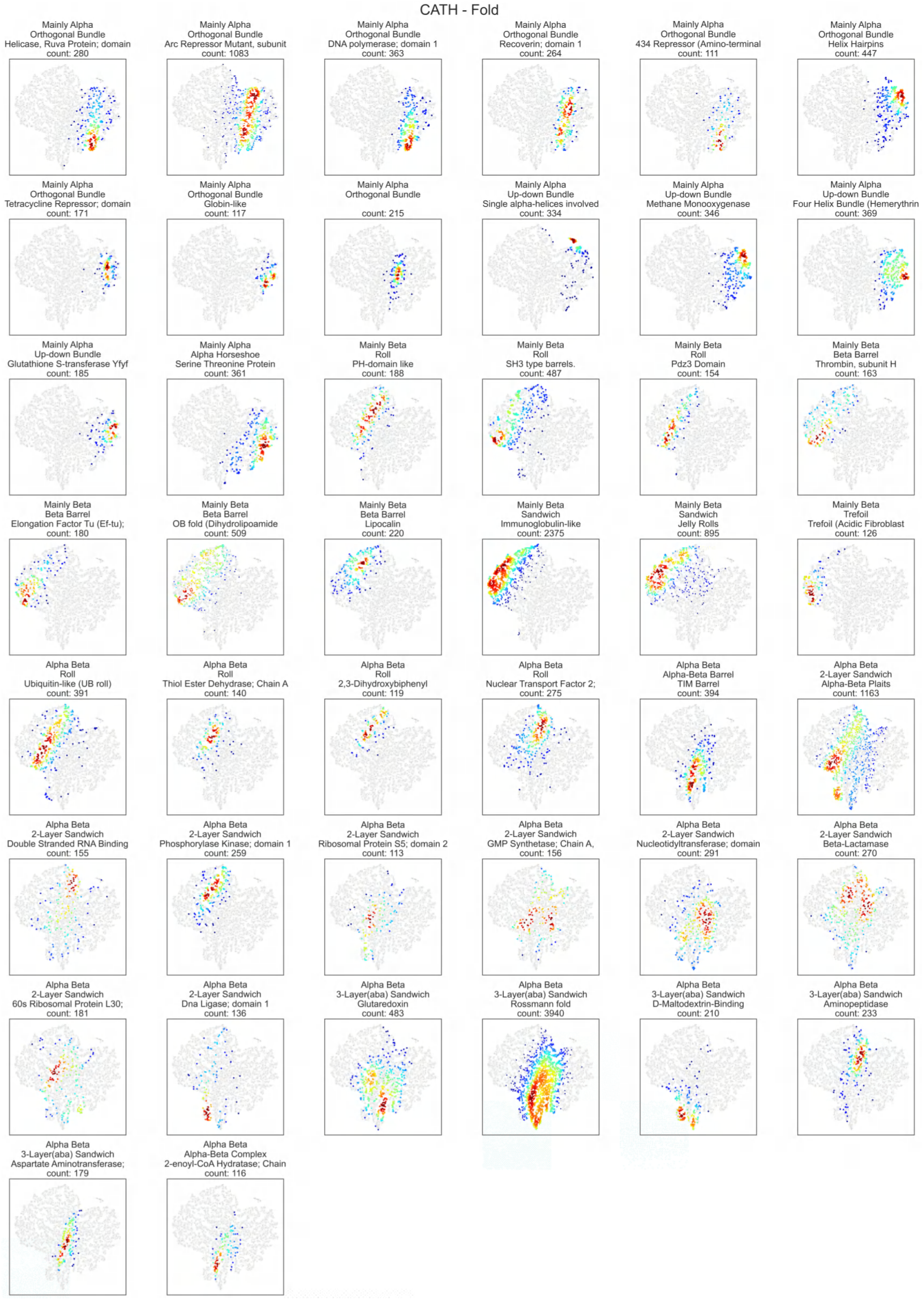
Density plot of CATH fold groups on the latent space. Each subplot displays the density heatmap of a specific CATH fold group, where the samples are colored by the local density calculated using the Gaussian kernel density estimation. To simplify the visualization, only the 44 fold groups that have more than 110 proteins are shown.

**Figure S3.**
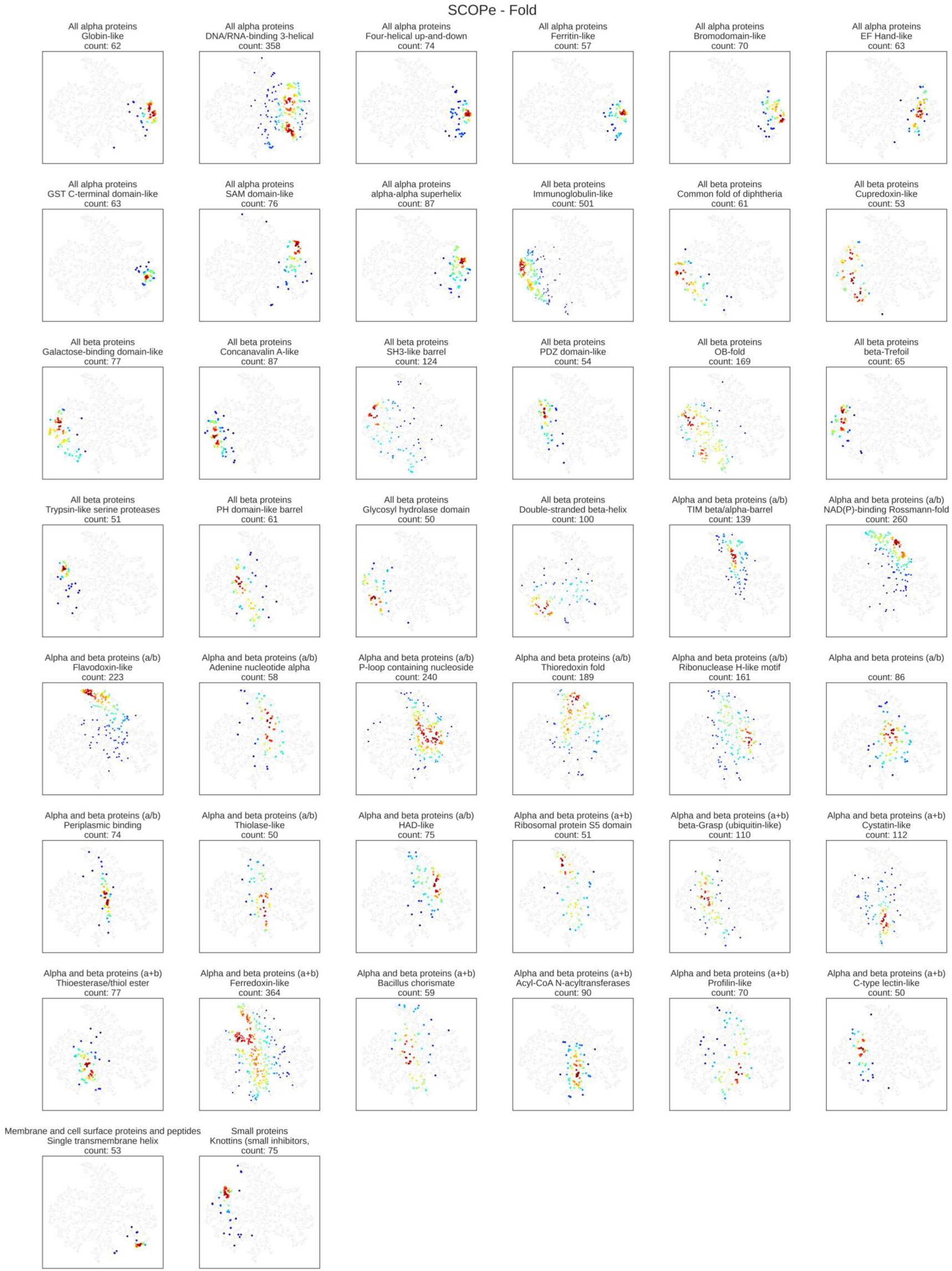
Density plot of SCOPe fold groups on the latent space. Each subplot displays the density heatmap of a specific SCOPe fold group, where the samples are colored by the local density calculated using the Gaussian kernel density estimation. To simplify the visualization, only the 44 fold groups that have more than 50 proteins are shown.

**Figure S4.**
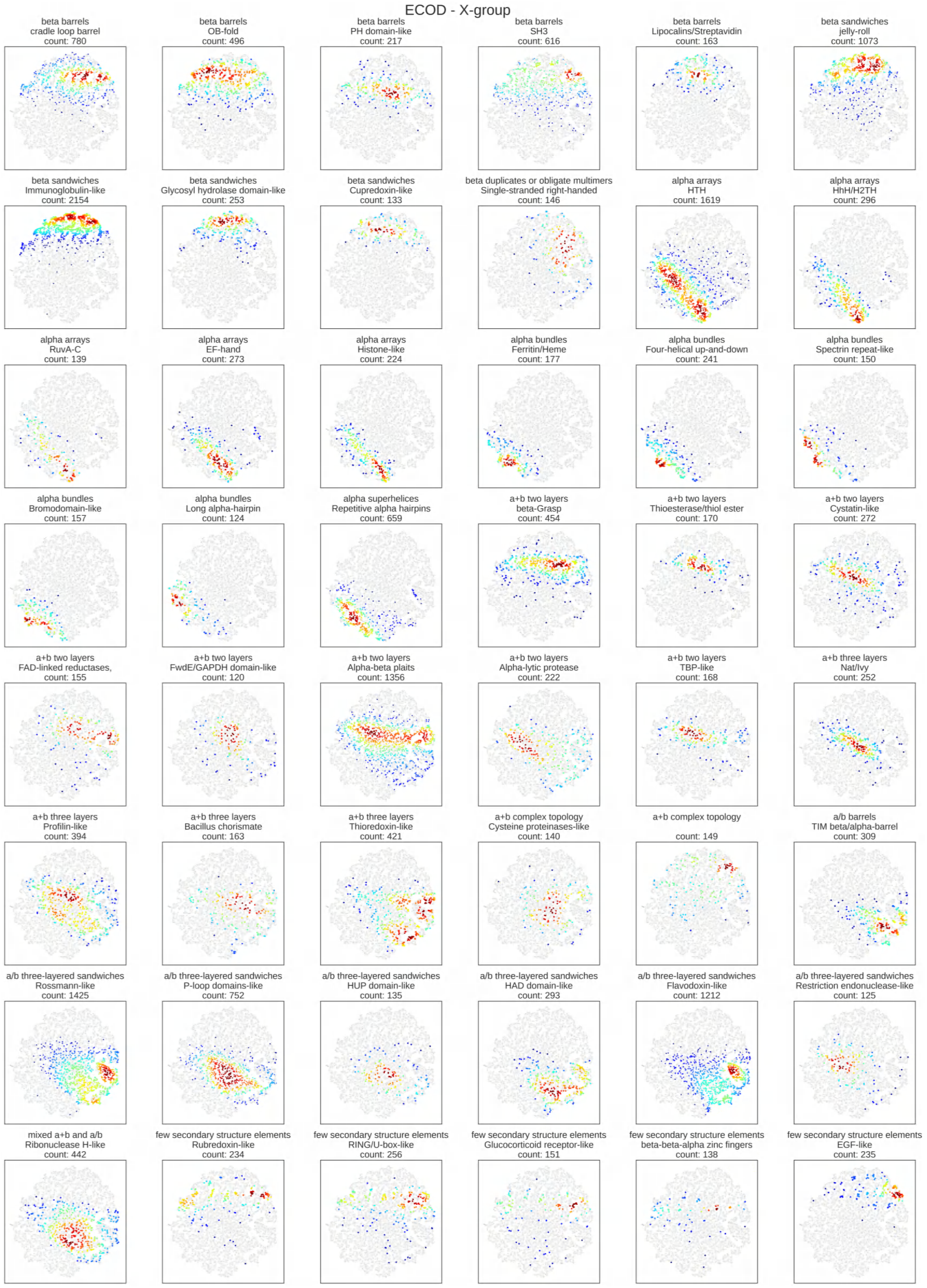
Density plot of ECOD fold groups on the latent space. Each subplot displays the density heatmap of a specific ECOD X-group, where the samples are colored by the local density calculated using the Gaussian kernel density estimation. To simplify the visualization, only the 48 groups that have more than 110 proteins and the recorded group names are shown.

#### 2 Wasserstein distance of latent distribution across CATH architectures

**Figure S5.**
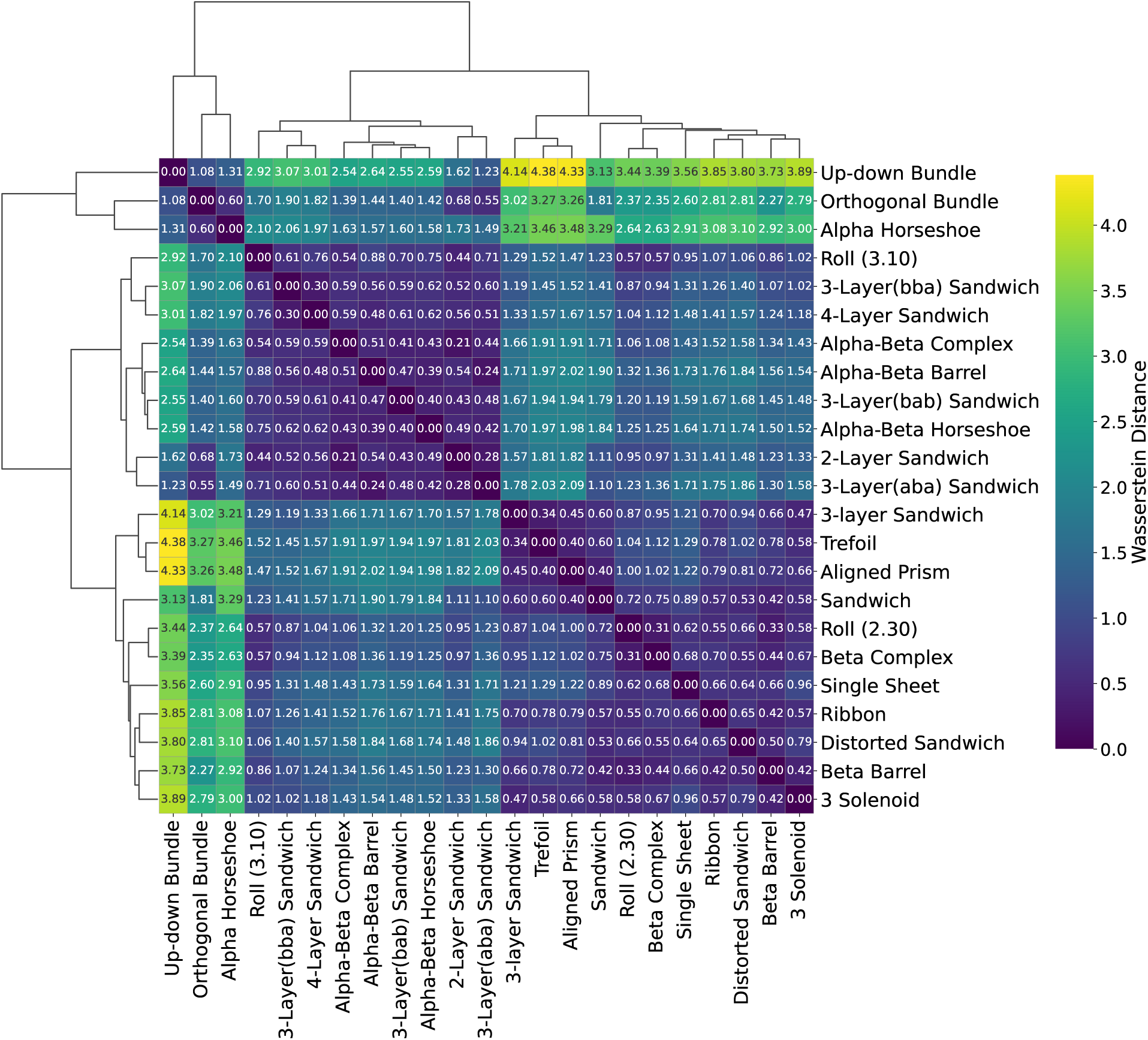
Clustermap of latent distribution distance across CATH architectures. The heatmap visualizes the pairwise Wasserstein distances (Equation 5) between latent distributions corresponding to different categories. The names of architectures are displayed at the bottom and right of the clustermap. Hierarchical clustering dendrograms positioned on the top and left illustrate the grouping of categories based on their distributional similarities. Darker colors indicate smaller distances, representing greater similarity between their latent distributions, while lighter colors indicate larger distances, suggesting greater dissimilarity. Annotations within individual cells provide the exact EMD values.

#### 3 Filtered latent outliers from CATH Class and Architecture hierarchies

**Figure S6.**
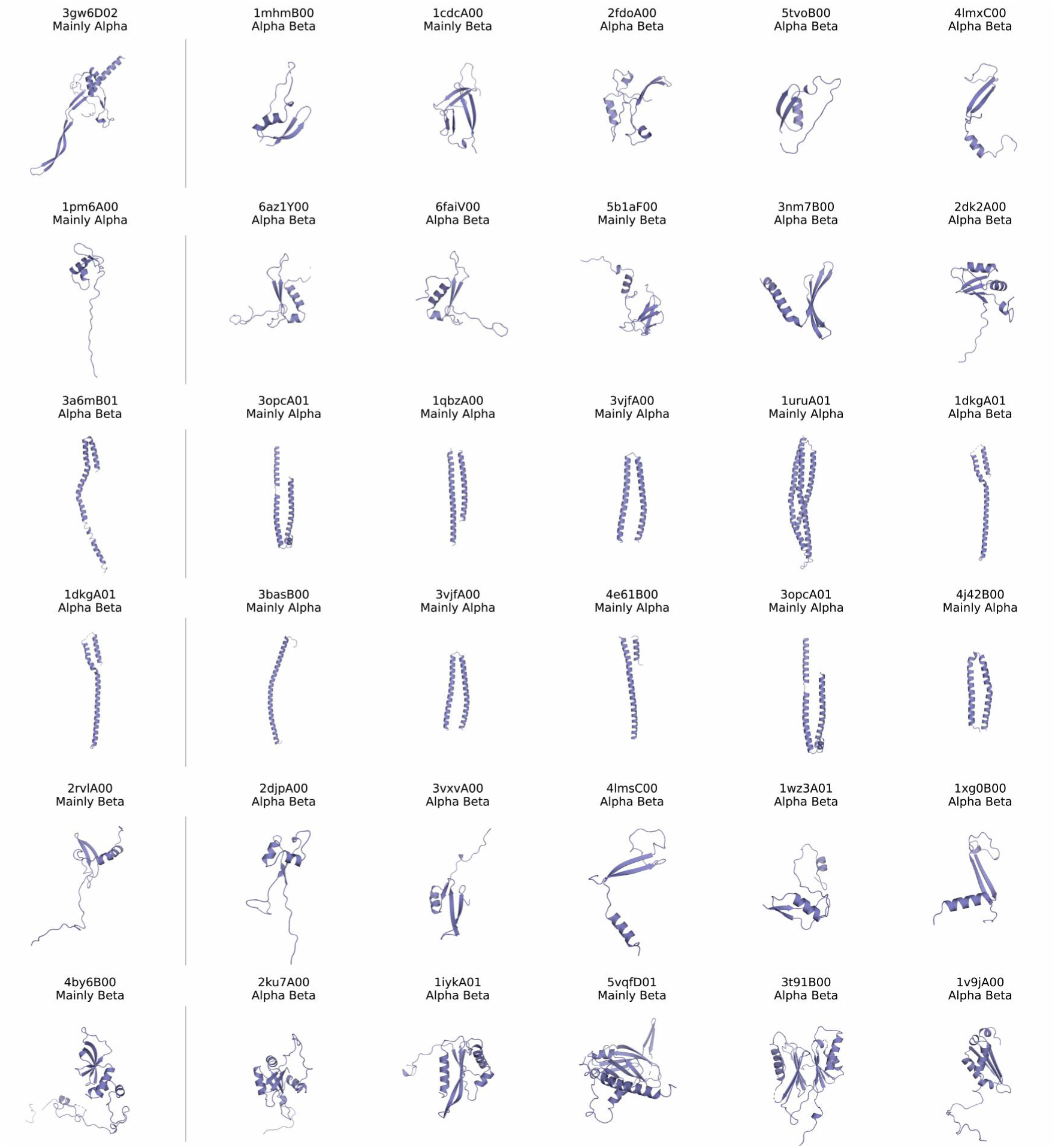
Filtered latent outliers at CATH Class hierarchy. This figure serves as a supplement to Figure 2E, displaying additional filtered samples of CATH Class hierarchy outliers. On the left side of each row, the annotation and structure of an outlier are presented, while its five closest neighbors in the latent space are shown on the right side. Each structure is labeled with its name and the associated CATH Class. The upper two rows, the middle two rows and the lower two rows show outliers of the Mainly-Alpha class, Alpha-beta class and Mainly-beta class, respectively.

**Figure S7.**
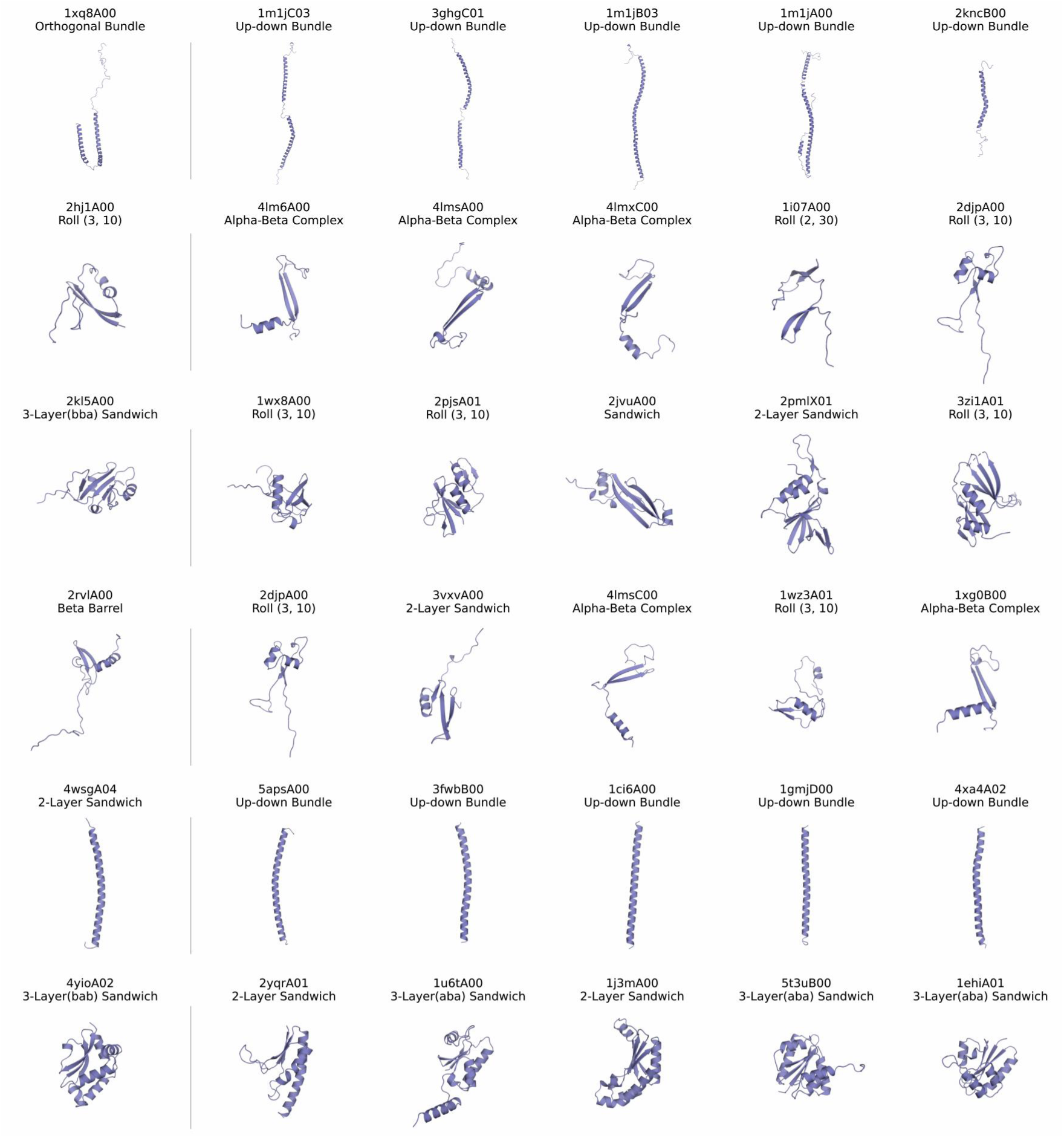
Filtered latent outliers at CATH Architecture hierarchy. This figure displays additional filtered samples of CATH Architecture hierarchy outliers. On the left side of each row, the annotation and structure of an outlier are presented, while its five closest neighbors in the latent space are shown on the right side. Each structure is labeled with its CATH ID and the associated CATH Architecture annotation.

#### 4 Additional results for benchmark on unconditional sampling

##### 4.1 Model parameter size and sampling efficiency

**Table S1.**
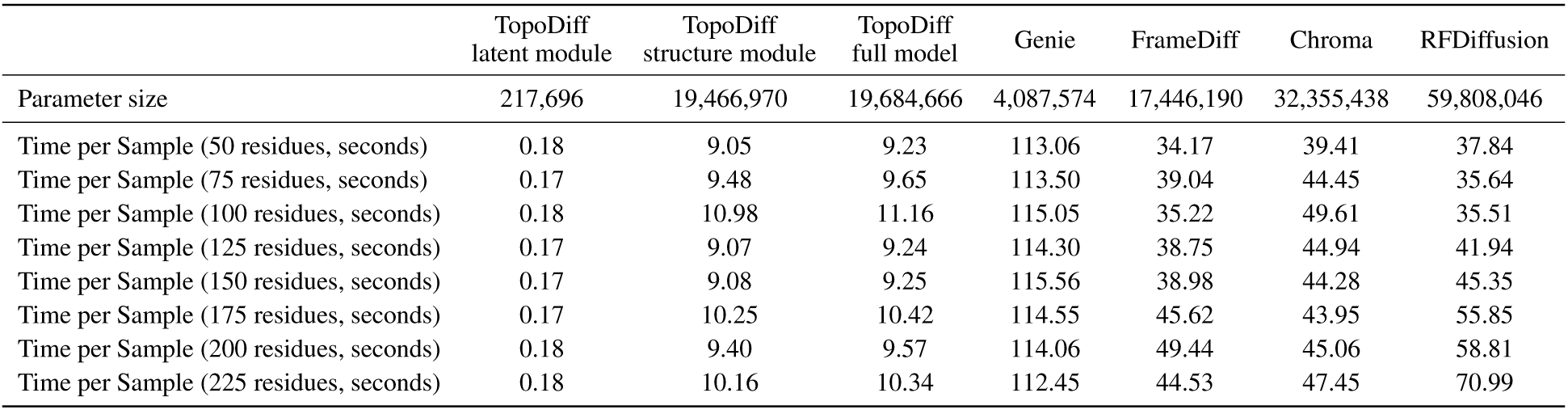
Comparison of parameter size and sampling efficiency.

To estimate the sampling efficiency of each model, we generated 10 backbones per length with each model and recorded the average time consumption. All models were sampled with a batch size of 1 on an NVIDIA Titan RTX GPU. For all models including TopoDiff, we conducted sampling using the provided scripts with minor modifications to exclude time for model initialization and to allow sampling with specified length and batch size. For TopoDiff, we also provided separate statistics for the two-stage sampling process (*i.e.* latent diffusion and structure diffusion) to offer a more detailed analysis.

##### 4.2 Topics on the coverage metric

We provide further discussion on the newly proposed coverage metric, particularly regarding the choice of hyperparameters and its potential for pinpointing underrepresented fold modes.

**Figure S8.**
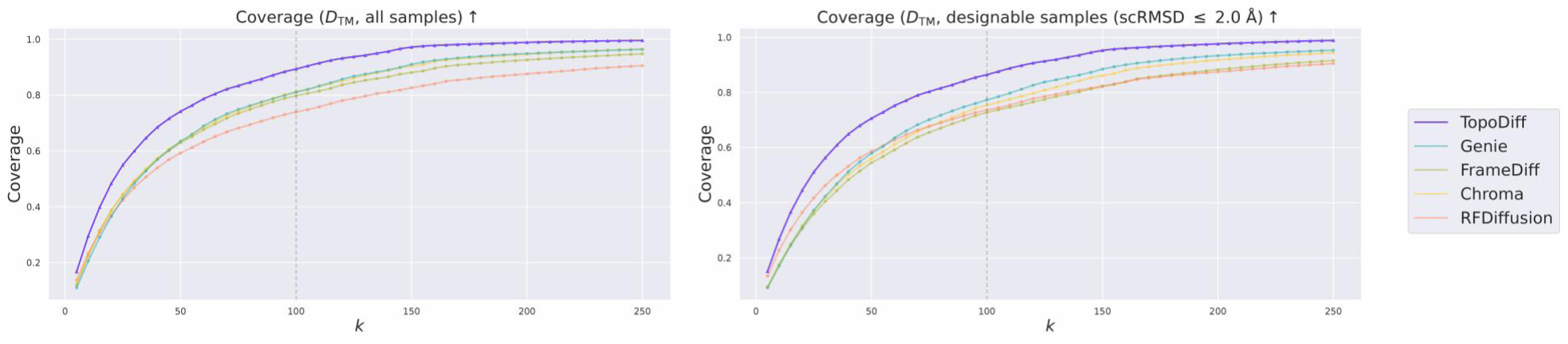
Coverage as a function of KNN hyperparameter *k*. This figure presents how the coverage changes as the KNN hyperparameter *k* is gradually increased. In the left subplot, coverage is computed using all samples generated by each method at each fixed length. In the right subplot, the computation focus on those generated samples with a designability cutoff (scRMSD *≤* 2 Å). In both subplots, *D*_TM_ (Equation 7) is used to define the distance between two samples.

As detailed in the “Methods” section, a hyperparameter *k* is used to define the radius of neighborhood hypersphere among real samples (*i.e.* reference natural proteins) to build the nearest-neighbor manifolds (Equation 6). A real sample is considered covered only if at least one generated sample falls within this hypersphere. Thus, a higher value of *k* results in a larger hypersphere and, consequently, a higher coverage estimate, and vice versa. In Supplementary Figure S8, we plotted the variation of the average coverage performance of each method as a function of the hyperparamter *k*. While the choice of *k* affects the exact coverage values, the relative ranking of the methods remains largely unchanged, and TopoDiff consistently demonstrates a clear advantage over the others, when measured both across all generated samples and across those exceeding a designability cutoff. We reported the coverage at the choice of *k* = 100 for all the other experiments.

**Figure S9.**
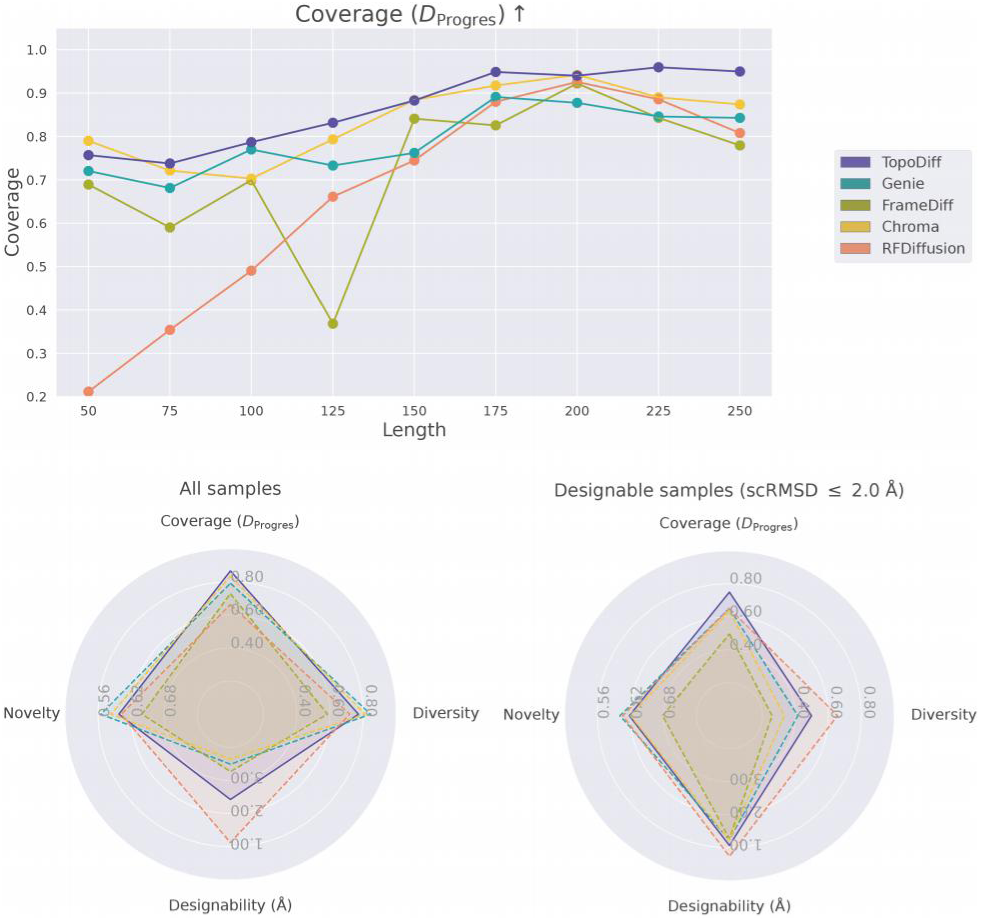
Comparison of coverage for unconditional sampling with *D*_Progres_ as distance definition. This figure presents the coverage metric recalculated using *D*_Progres_ as the distance definition, instead of *D*_TM_. The upper panel shows the coverage at each sampled length, and the lower panel are radar plots with new coverage implementation. All other settings remain identical to those in the benchmark shown in Figure 3.

Another aspect of flexibility in the implementation is the choice of distance function *D*(*·, ·*) to measure the distance/dissimilarity between structures. In the benchmark shown in Figures 3, we always used *D*_TM_ (Equation 7) to define the distance. Since the computation of pairwise distances between real samples and generated samples can be computationally expensive with this implementation, we also tested an alternative distance definition *D*_Progres_ (Equation 8), which employs a third-party model to encode all structures into vector representations on a hypersphere, enabling parallel computation of cosine similarity. In Supplementary Figure S9, we presented the benchmark results for the unconditional sampling when this alternative distance definition is used. TopoDiff consistently exhibits advantage in the mode coverage compared to the other methods.

**Figure S10.**
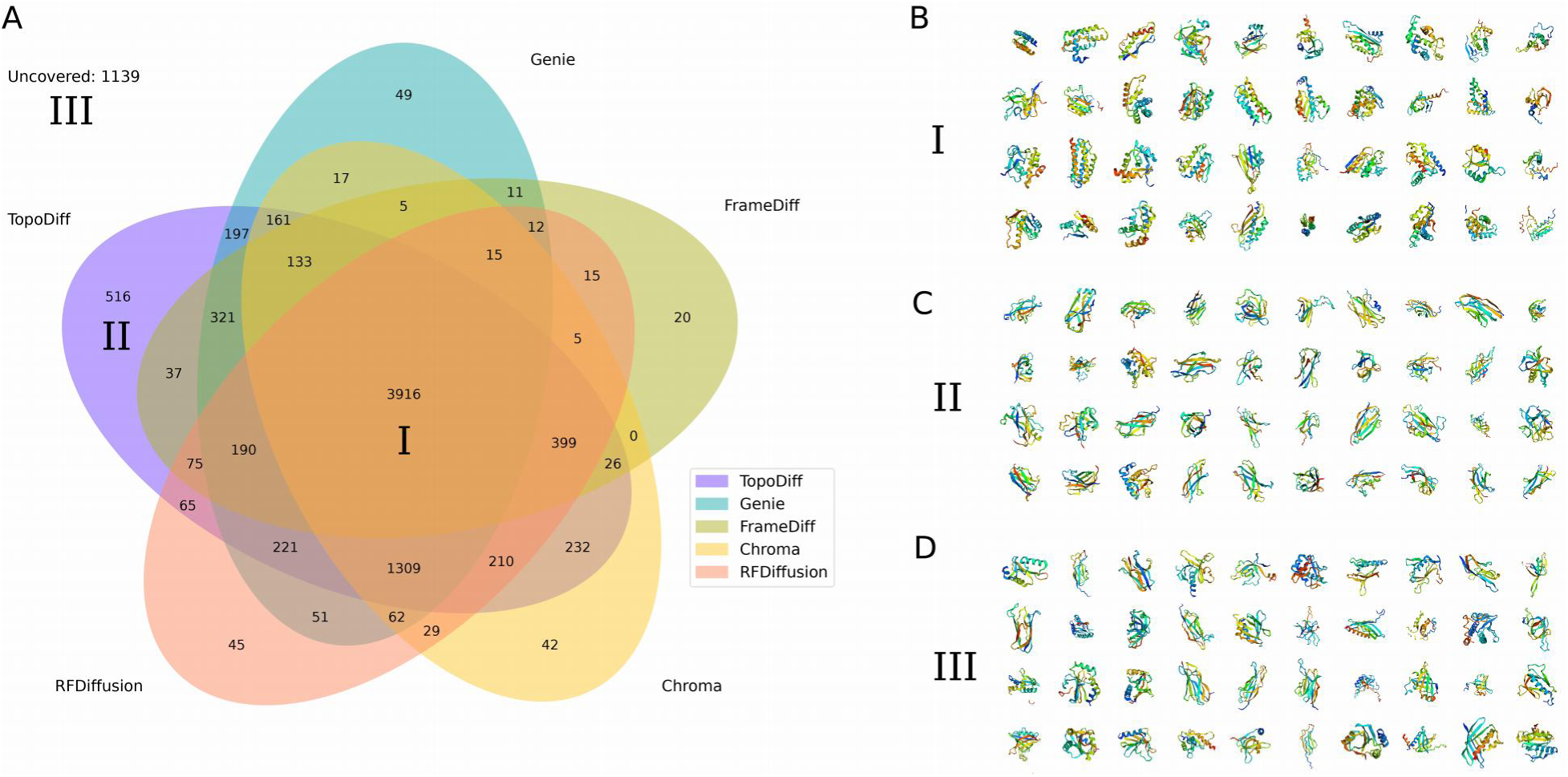
Analysis of fold coverage across models using sample-wise binary indicators (length 125). **(A)** Venn diagram showing the overlap in the coverage of natural protein folds (length range: 100 ∼ 150) among the five models under comparison. The number of natural fold modes in each group is displayed within the respective region. Three regions are highlighted for further visual inspection: folds covered by all five models (region I), folds uniquely covered by TopoDiff (region II), and folds not covered by any of the models (region III). **(B)** Representative samples from region I (folds covered by all models). **(C)** Representative samples from region II (folds uniquely covered by TopoDiff). **(D)** Representative samples from region III (uncovered folds).

By definition, the coverage is computed by averaging the binary indicators for each real sample denoting it is covered by generated samples or not. Therefore, besides the final coverage value, we could also analyze the sample-wise binary indicator to gain a deeper understanding of the covered or uncovered fold modes. Here, we illustrate this analysis using generated samples with the length of 125. First, we presented a Venn diagram comparing the five models under evaluation to visualize the overlapping coverage of natural folds (Supplementary Figure S10A). Out of the 9,525 natural folds within the length range of [100, 150], 3,916 folds are commonly covered by all of the 5 models (region I). TopoDiff stands out by covering a total of 8,008 folds, with a unique coverage of 516 folds (region II). Additionally, there are also 1,139 natural folds that are not considered as covered by any of the five models (region III). To gain a more intuitive understanding of the characteristics of the samples in these regions (regions I, II and III), we randomly selected 40 samples from each region and visualized them at the right panel. For the commonly covered natural folds (Supplementary Figure S10B), most of them consist of a high proportion of alpha helices and are generally well-packed into a globular shape. Further examination of the samples uniquely covered by TopoDiff (Supplementary Figure S10C) reveals that our model could cover a large number of natural folds characterized by the mainly beta composition. The folds commonly left uncovered by all models (Supplementary Figure S10D) are largely beta proteins as well, often displaying more complex topologies or irregular global shapes. Overall, our result demonstrates that there is still room for improvement in the current generative models on the successful modeling of proteins with complex topology or shape. While our model fails to approach the ultimate perfection, the improvement is already visually intuitive and the development of the new coverage metric along with the sample-wise binary indicators could help us address underrepresented folds and give direction for future improvement.

#### 5 Additional results for controllable generation

**Figure S11.**
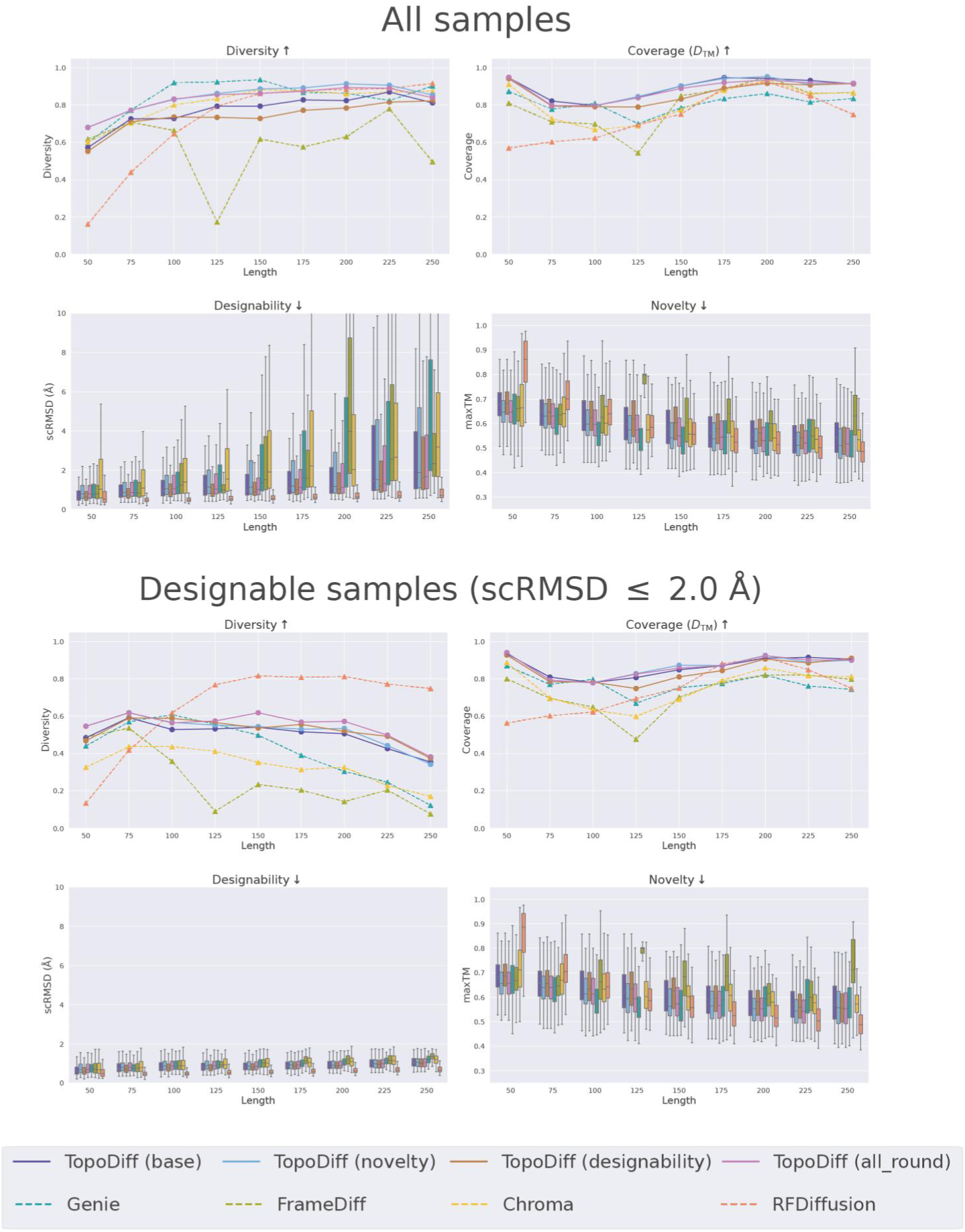
Additional result on controlling sampling distribution by latent space rejection sampling. This figure supplements Figure 4B. The detailed performance at each sampled length of four model variants of TopoDiff achieved by tuning the threshold of latent-based rejection sampling, along with other benchmarked methods, are displayed. In the upper panel, each metric is calculated using all samples generated at individual lengths. In the lower panel, only the highly designable samples with scRMSD *≤* 2 Å are included in the metric computations.

**Figure S12.**
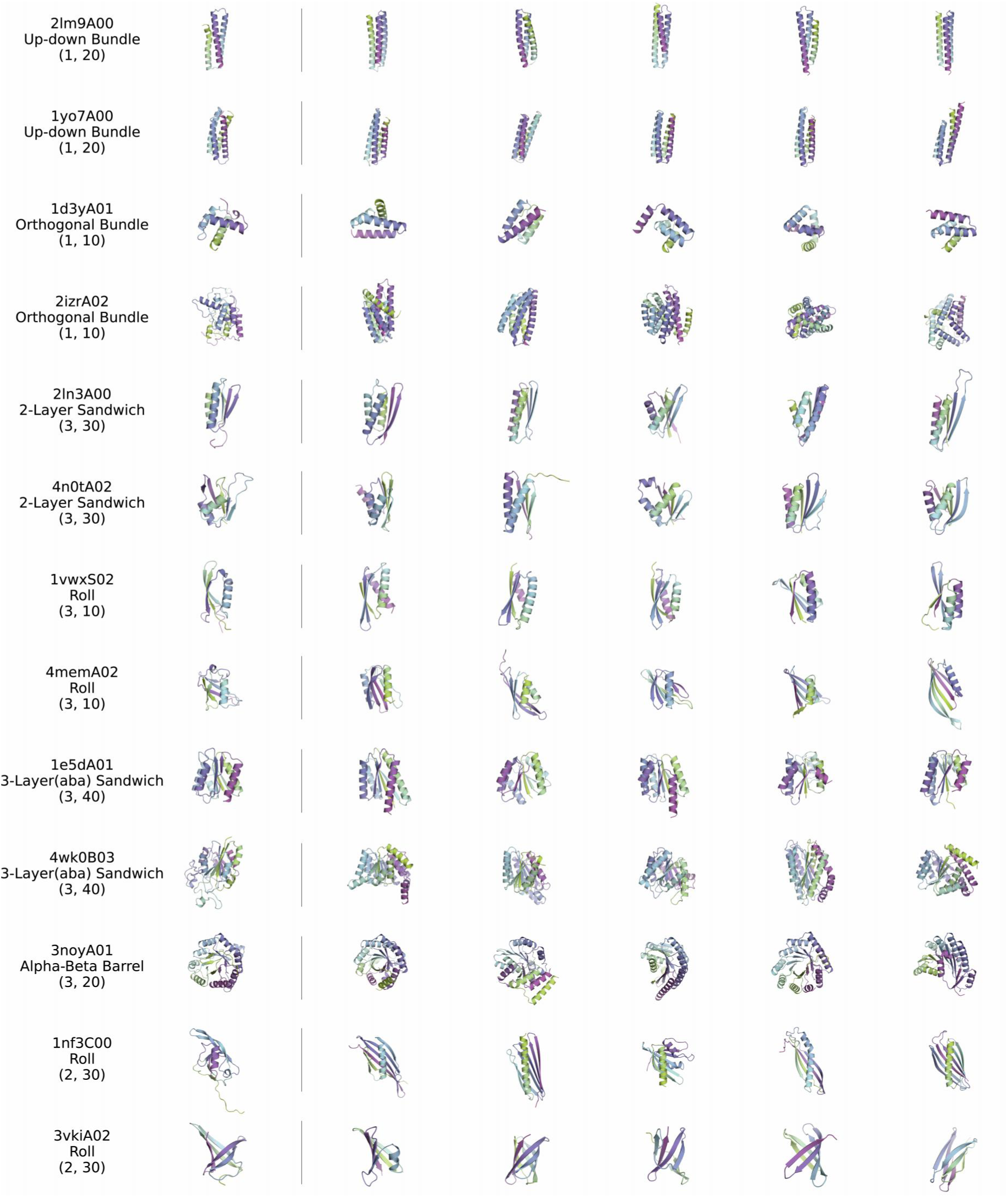
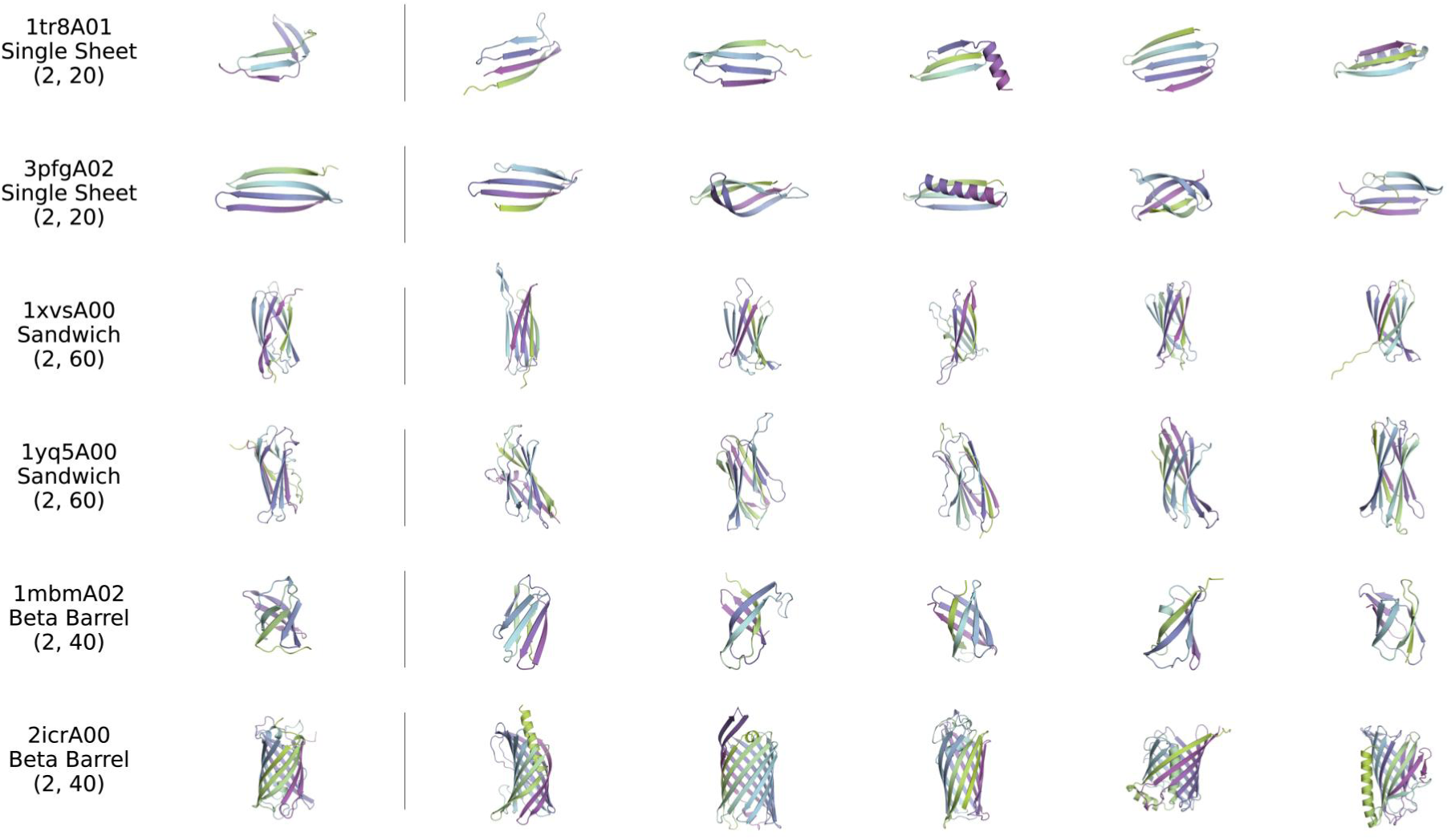
Sampling proteins with similar global geometry. This figure supplements Figure 4C. For each representative protein selected from the CATH-60 dataset, its structures is encoded by the structure encoder, and the resultant latent code is used to sample 5 new structures. On the left side of each row, the annotation and structure of the reference structure are presented, and the 5 randomly sampled proteins without cherry-picking are shown on the right side. Each reference structure is labeled with its CATH ID and the associated CATH Architecture annotation.

**Figure S13.**
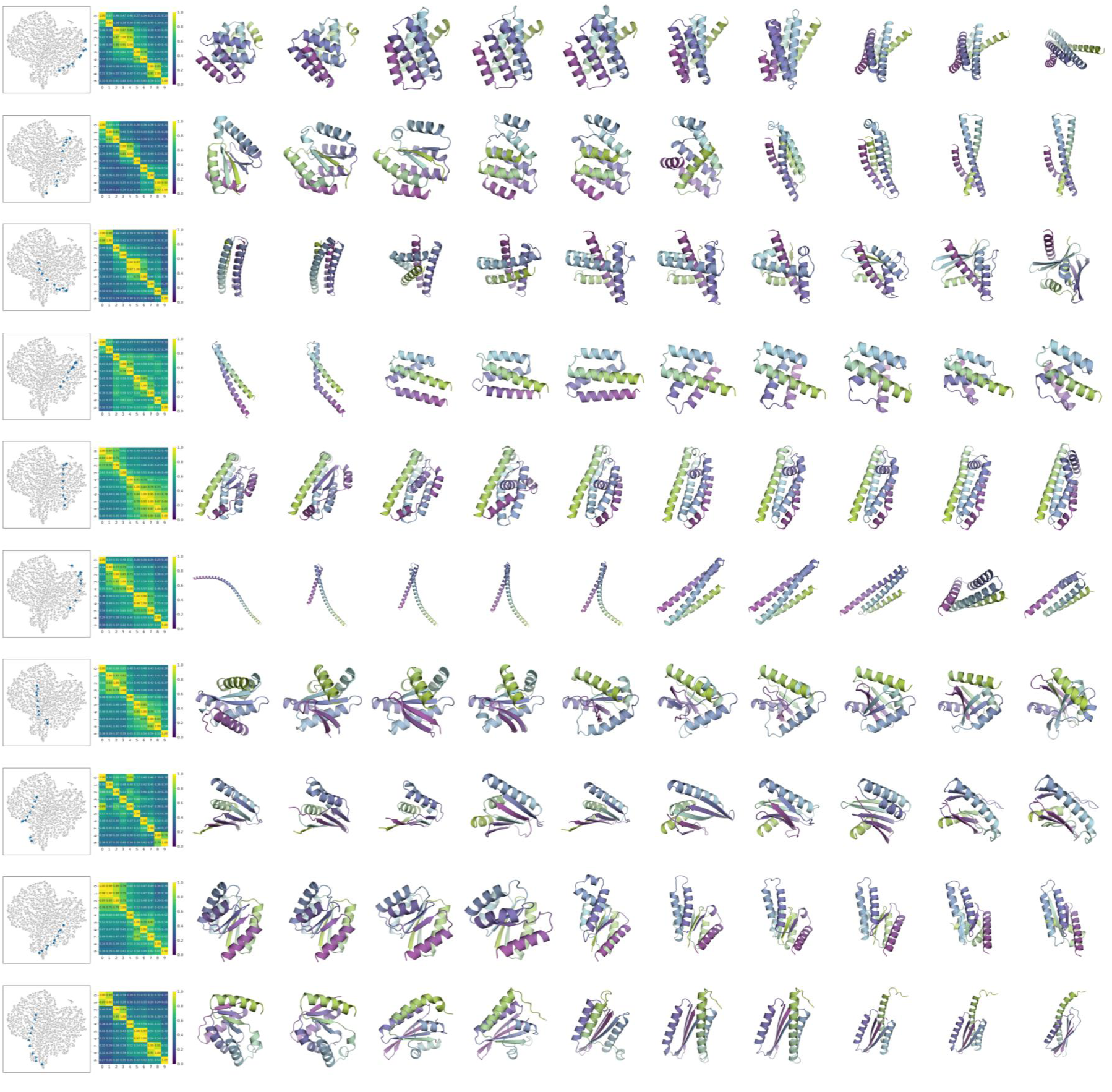
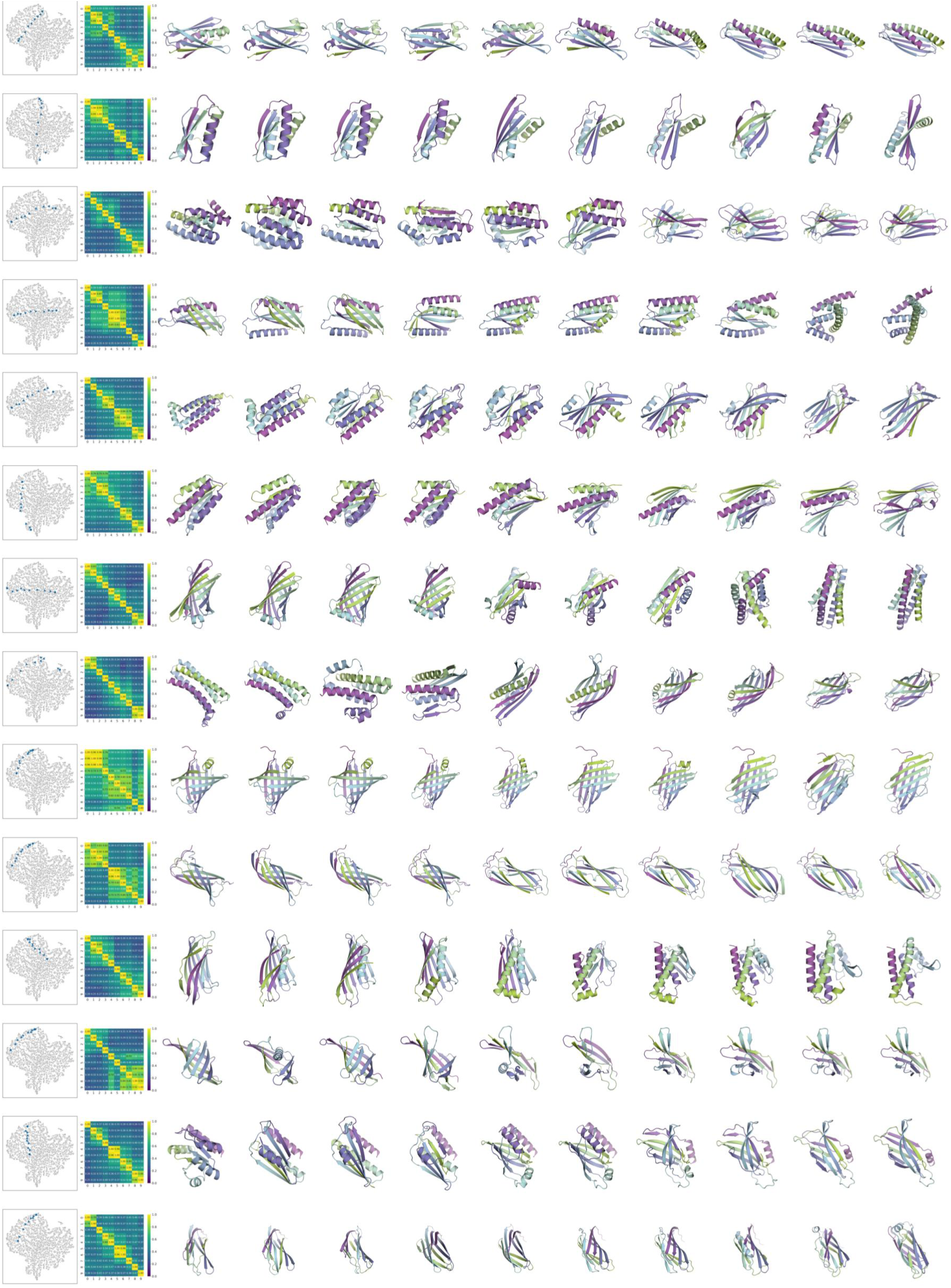
Additional examples for sampling proteins with latent code interpolation. This figure supplements Figure 4D, presenting more examples of protein sampling with the latent code interpolation. We selected pairs of latent codes across the latent distribution to demonstrate the continuity of the latent space and the smooth transitions between sampled structures. Each row corresponds to an individual interpolation result. The leftmost panel shows the linear interpolation trajectory between two latent codes, projected onto the t-SNE dimension-reduced space, with the direction indicated by the intermediate triangles. The second panel shows the pairwise TM-score matrix of the ten sampled structures, with their detailed structures visualized on the right.

#### 6 Additional results for experimental validation

##### 6.1 Summary of the selected candidates

**Figure S14.**
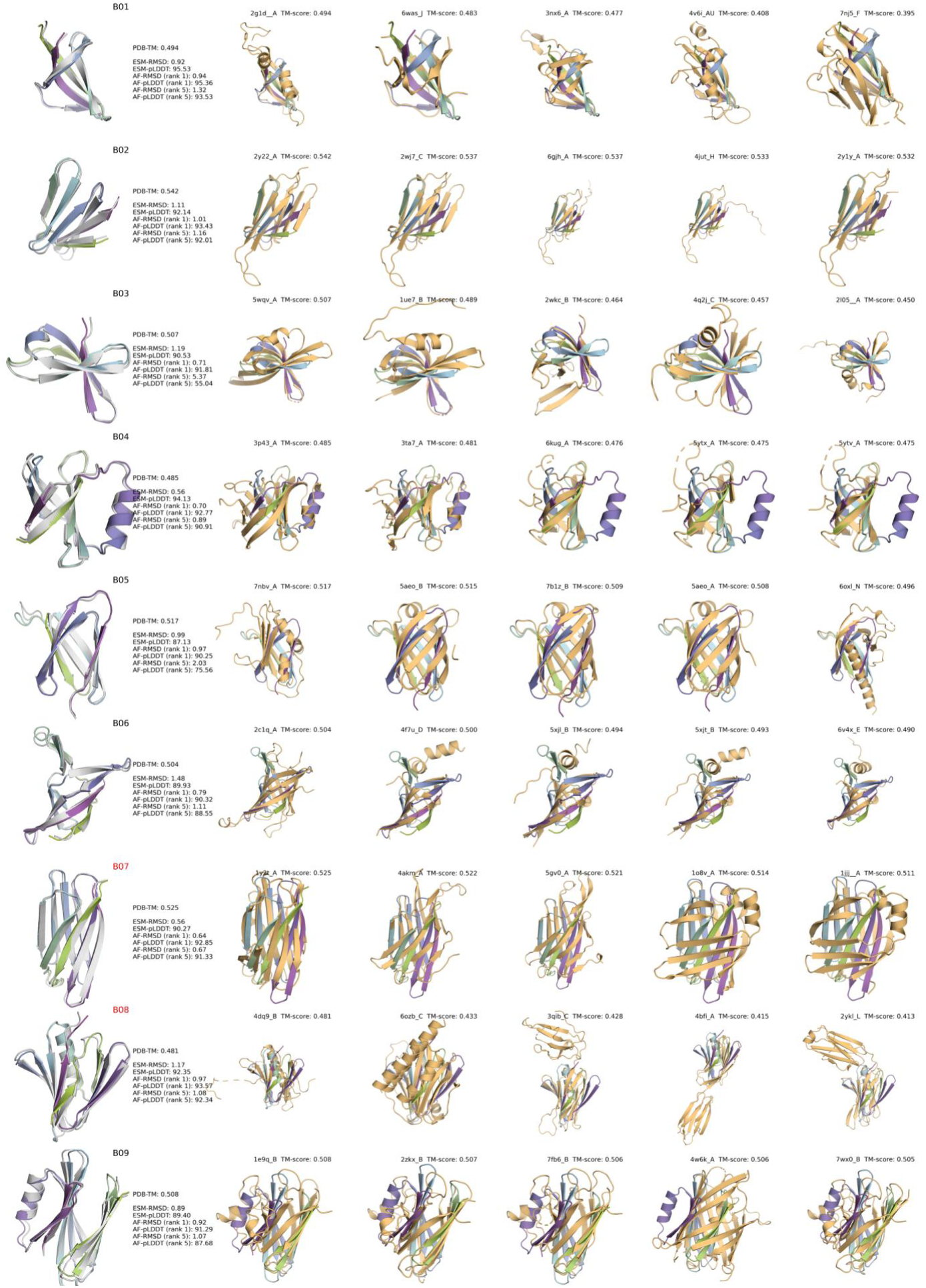
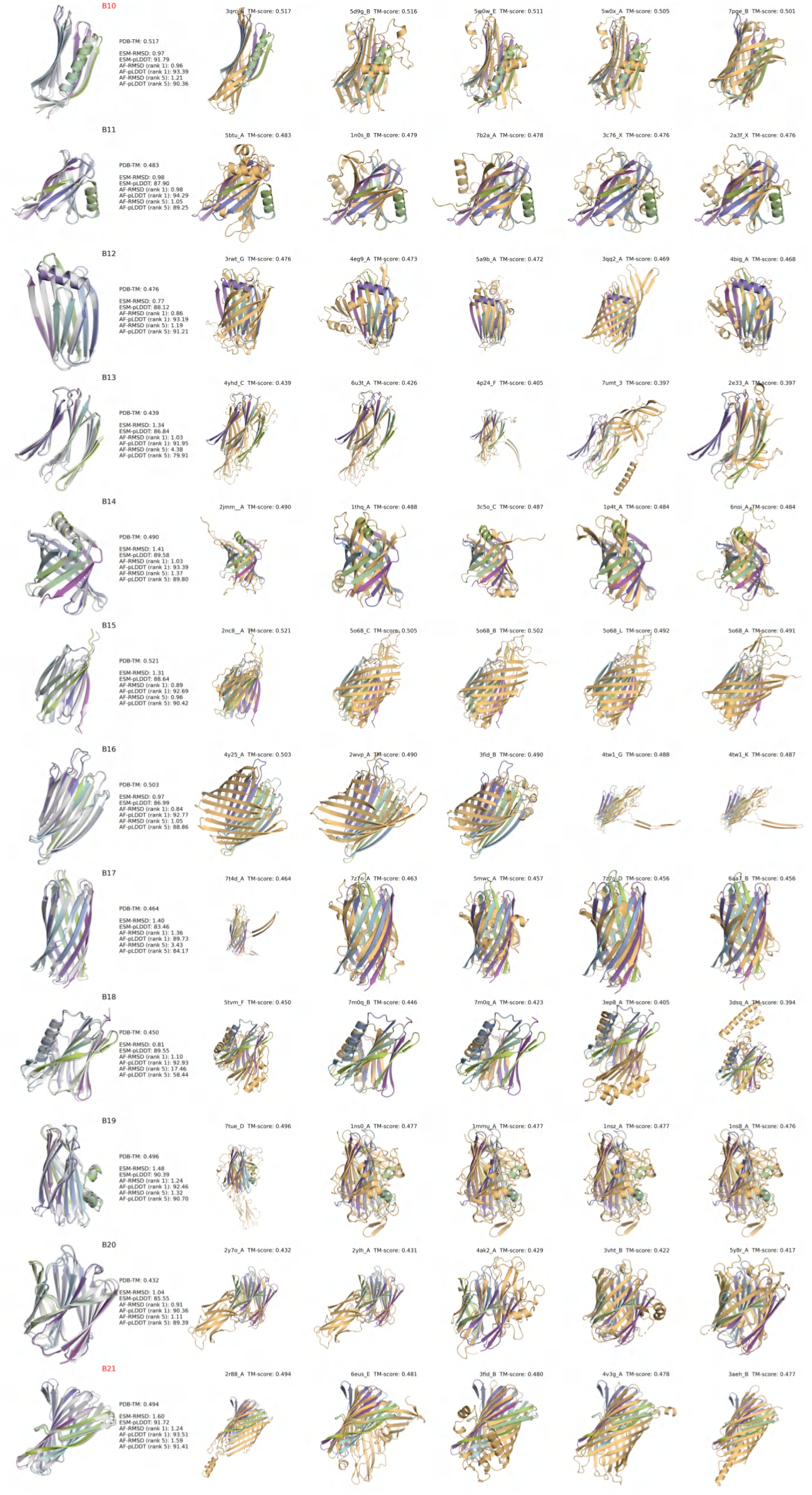
Visualization of selected candidates and their closest structures in PDB. Each row represents a candidate design. In the leftmost panel, the design backbone is displayed along with its corresponding metrics: PDB-TM (maximal TM-score relative to PDB structures), ESM-RMSD (scRMSD predicted by ESMFold), ESM-pLDDT (confidence score by ESMFold), and AF-RMSD & AF-pLDDT (similar metrics predicted by AlphaFold2). Rankings (*e.g.*, 1*^st^*, 5*^th^*) indicate the prediction order among AlphaFold2’s five models, based on AF-RMSD. On the right, the five closest structures from PDB identified by Foldseek search are shown along with the corresponding query TM-scores.

##### 6.2 Summary of experimental results

**Table S2.** Summary of experimentally validated designs.

| Sequence Length | Scaffold ID | Design ID | *E. coli* Expression | Solubility | SEC | β-sheet-rich CD Spectrum | $T_m$ (°C) |
| --- | --- | --- | --- | --- | --- | --- | --- |
| 50 | b_050_0 | B01 | no | no |  |  |  |
|  | b_050_2 | B02 | no | no |  |  |  |
|  | b_050_4 | B03 | no | no |  |  |  |
| 75 | b_075_0 | B04 | no | no |  |  |  |
|  | b_075_1 | B05 | no | no |  |  |  |
|  | b_075_2 | B06 | no | no |  |  |  |
| 100 | b_100_0 | B07 | yes | yes | monomer | yes | > 95 |
| | b_100_1 | B08 | yes | yes | monomer & oligomer | yes | ≈ 80.4 |
|  | b_100_2 | B09 | yes | yes | oligomer | no |  |
| 125 | b_125_0 | B10 | yes | yes | monomer | yes | > 95 |
|  | b_125_1 | B11 | no | no |  |  |  |
|  | b_125_2 | B12 | yes | yes | oligomer | yes |  |
| 150 | b_150_0 | B13 | no | no |  |  |  |
|  | b_150_1 | B14 | no | no |  |  |  |
|  | b_150_2 | B15 | yes | yes | oligomer | no |  |
| 175 | b_175_0 | B16 | yes | yes | oligomer | yes |  |
|  | b_175_1 | B17 | no | no |  |  |  |
|  | b_175_2 | B18 | no | no |  |  |  |
| 200 | b_200_0 | B19 | yes | yes | oligomer | no |  |
|  | b_200_1 | B20 | no | no |  |  |  |
| | b_200_2 | B21 | yes | yes | monomer & oligomer | yes | ≈ 65.2 |

$T_m$ refers to the melting temperature. All designs were expressed in *E. coli* and purified by nickel-affinity chromatography, followed by SEC and CD measurements. Soluble monomeric designs were tested for melting temperature.

##### 6.3 Protein expression and purification

**Figure S15.**
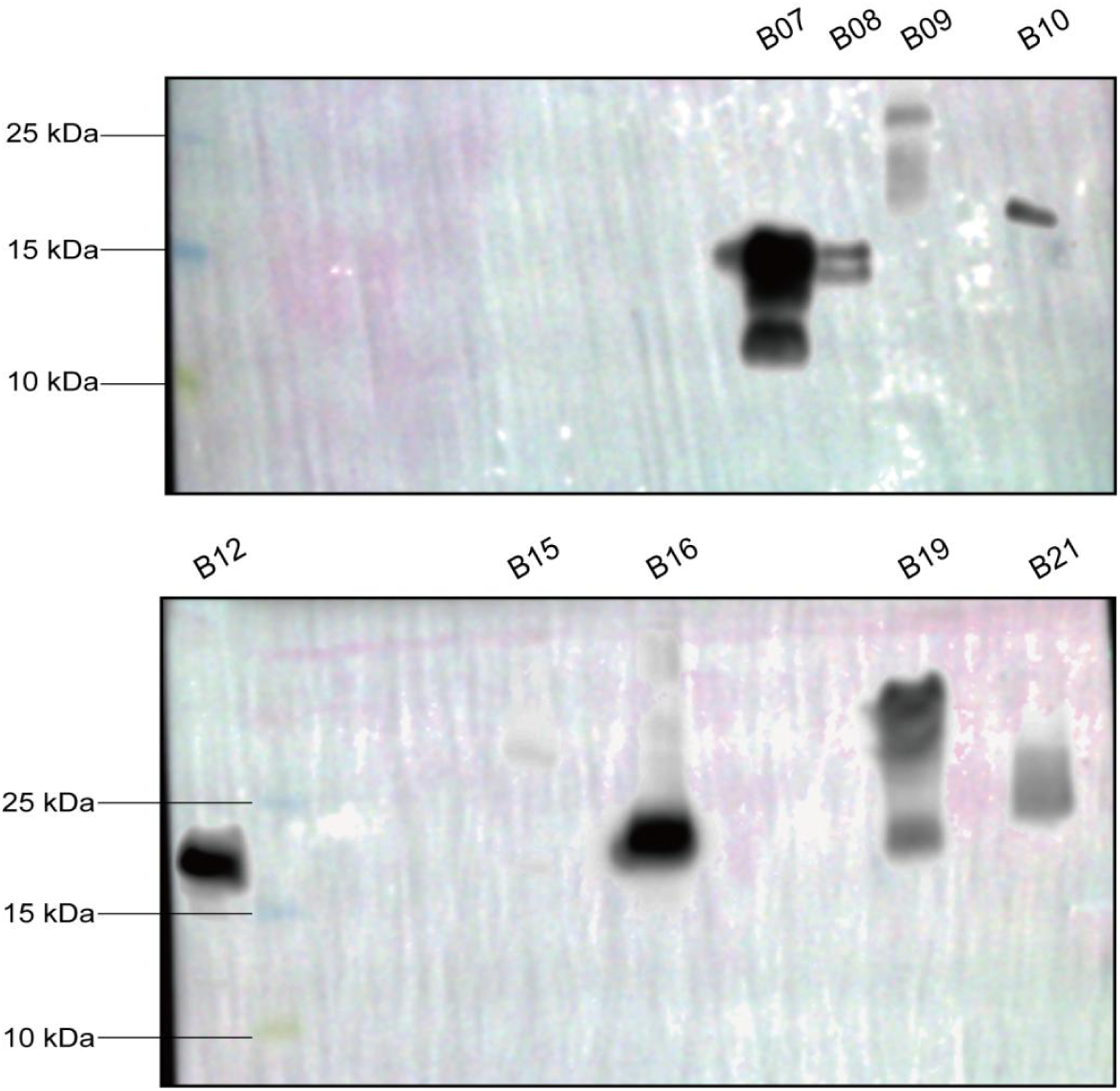
Western blot detecting the expression of selected designs in *E. coli*. Nine out of twenty-one designs show a positive signal against the His-tag, and five out of nine expressed designs have a single band with the correct molecular weight.

**Figure S16.**
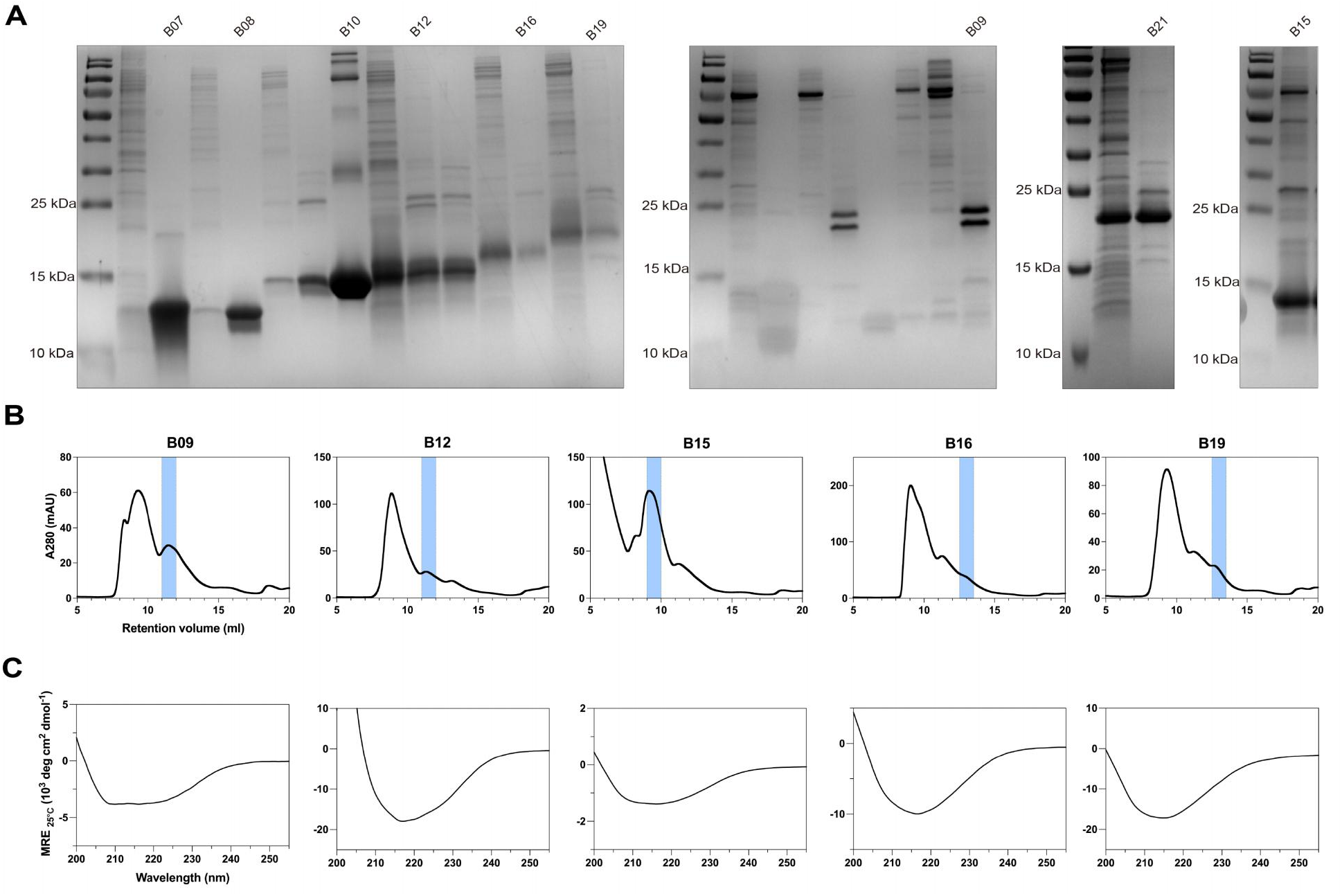
Experimental validation of the remaining expressed designs. **(A)** SDS-PAGE of purified protein fraction from SEC. The annotated bands, identified as monomeric designs, were utilized for CD measurement. **(B)** The SEC profiles with the monomeric peaks highlighted in blue. **(C)** The CD spectra of the purified proteins at 25*^◦^*C.

##### 6.4 X-ray structure determination

**Figure S17.**
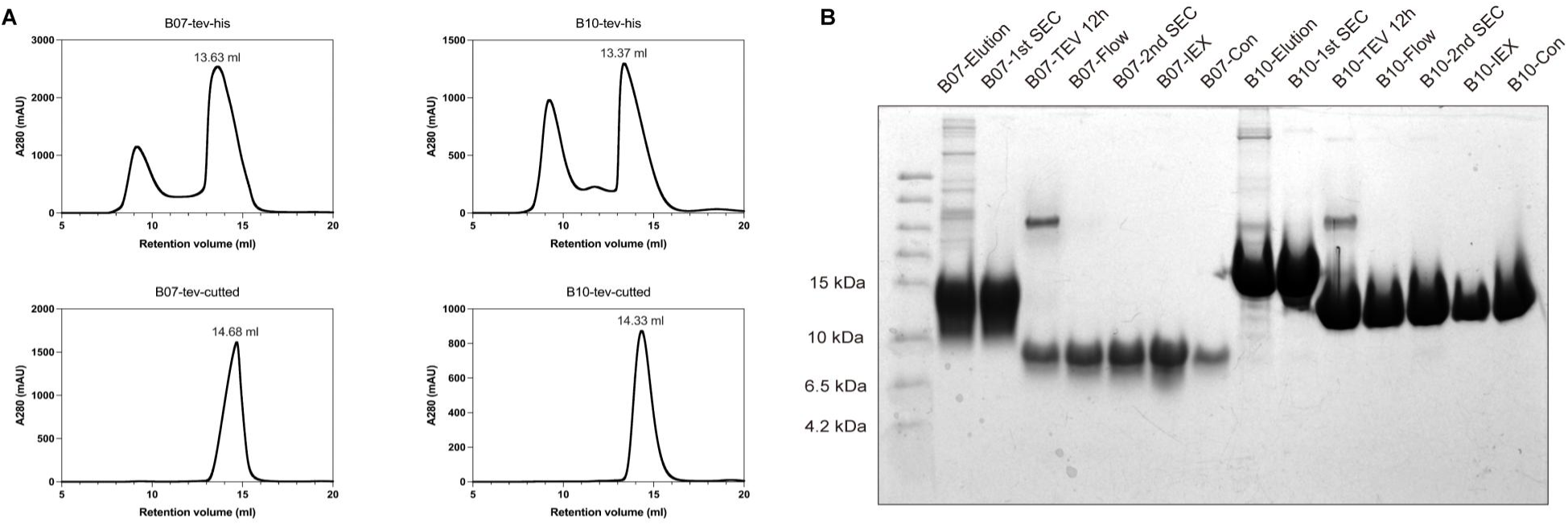
Preparation of crystallization sample for X-ray structural determination. The monomer designs B07 and B10 were modified to include a TEV cleavage site to remove the His-tag for crystallization. Purified samples were subjected to SEC, incubated with His-tagged HyperTEV protease at 4*^◦^*C for 12 hours, and flowed through Ni-NTA beads to remove His-tag and HyperTEV, followed by a second SEC and concentration. **(A)** The SEC profiles of the designs before and after the removal of His-tag. The monomeric peak shifted after cleavage. **(B)** Tricine-SDS-PAGE analysis of each component during the sample preparation process.

**Table S3.** Crystallographic data for design B10.

|  | B10 (monoclinic) | B10 (orthorhombic) |
| --- | --- | --- |
| Wavelength (Å) | 0.9789 | 0.9789 |
| Resolution range (Å) | 28.71 - 1.8 (1.865 - 1.8) | 29.58 - 1.702 (1.763 - 1.702) |
| Space group | C 2 | P 212121 |
| Unit cell | 70.886 47.68 38.93 90 97.956 90 | 80.879 85.163 94.642 90 90 90 |
| Unique reflections | 11988 | 71785 (6780) |
| Completeness (%) | 99.52 (98.59) | 99.40 (95.75) |
| Redundancy | 5.9 (4.3) | 10.0 (7.4) |
| Mean I/sigma(I) | 51.09 (1.05) | 36.56 (1.63) |
| Wilson B-factor | 14.42 | 22.79 |
| R-merge | 5.4 (13.4) | 8.5 (92.2) |
| CC1/2 | 100.2 (98.2) | 99.8 (72.9) |
| Reflections used in the refinement | 11974 (1188) | 71778 (6777) |
| Reflections used for R-free | 605 (57) | 2007 (187) |
| R-work | 0.174 | 0.1775 |
| R-free | 0.1984 | 0.2075 |
| Number of non-hydrogen atoms | 1164 | 4549 |
| macromolecules | 1007 | 4117 |
| ligands | 0 | 0 |
| Solvent | 157 | 432 |
| RMS (bonds) | 0.007 | 0.016 |
| RMS (angles) | 1.01 | 1.59 |
| Ramachandran favored (%) | 99.19 | 98.98 |
| Ramachandran allowed (%) | 0.81 | 1.02 |
| Ramachandran outliers (%) | 0.00 | 0.00 |
| Rotamer outliers (%) | 0.88 | 1.28 |
| Clashscore | 1.91 | 1.49 |
| Average B-factor | 20.82 | 34.96 |
| macromolecules | 19.76 | 34.21 |
| ligands | 0 | 0 |
| solvent | 27.59 | 42.10 |

#### 7 Additional structural analysis on the successful designs

**Figure S18.**
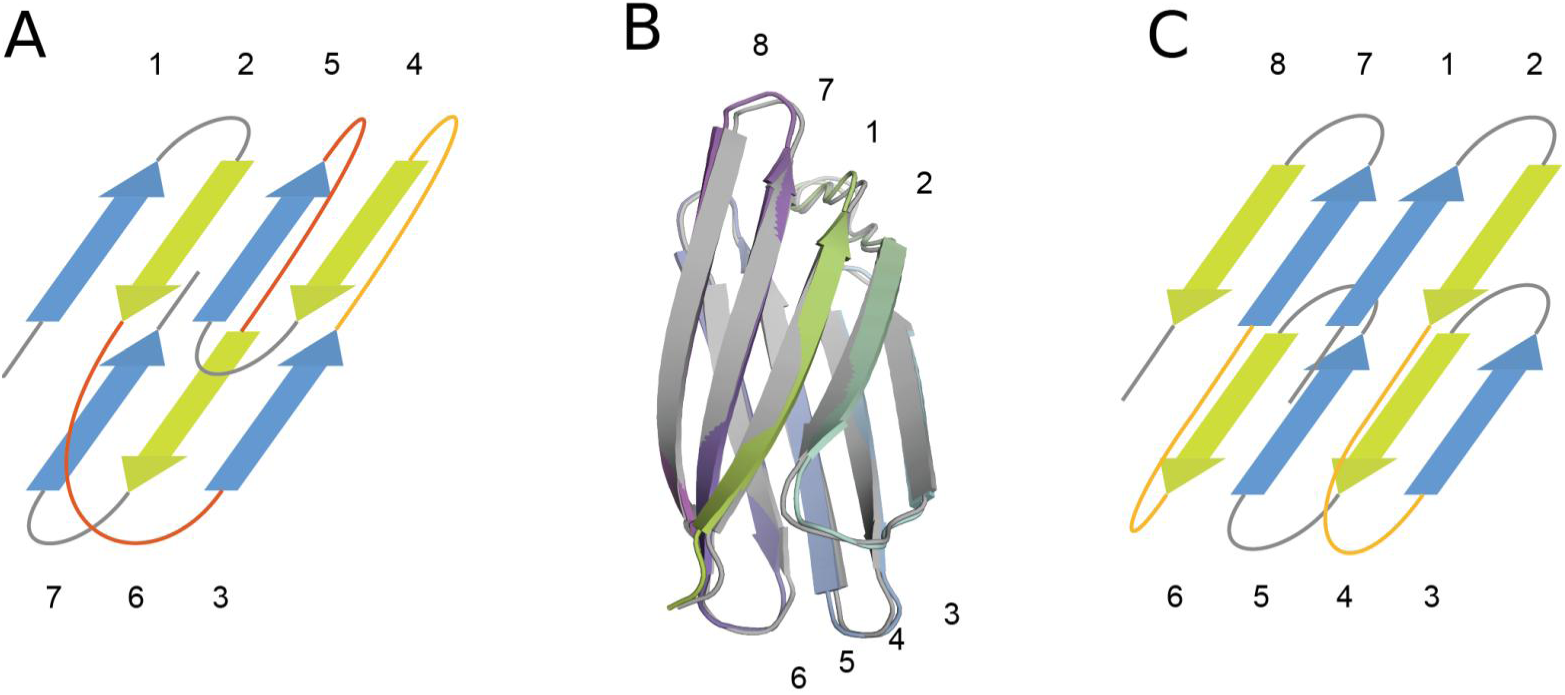
Structural analysis of design B07. **(A)** Example of a typical beta sandwich with the “interlock” arrangement (adapted from Kister et al.^1,2^). The two beta-arches (crossovers between two opposing *β*-sheets) colored in red form an “interlock” arrangement. Other beta-arches are colored in orange. **(B)** The designed backbone structure of B07 (colored) agrees with the AlphaFold2-predicted model (gray), with an RMSD of 0.64 Å. Each strand is numbered according to its sequential order. **(C)** The topology diagram of B07. The only two beta-arches are colored in orange.

As shown in Figure 5B&C, B10 is a 125-residue mainly-beta protein composed of 8 beta strands and 1 alpha helix, with the alpha helix inserted into the crossover region between the two beta sheets, creating a special triangular geometric arrangement. This unique geometry may create novel binding sites or interaction surfaces, potentially enabling the protein to interact with other molecules or ligands in a specific manner.

B07 is a 100-residue all-beta sandwich, with each beta sheet containing 4 strands, and the relative twist between the two sheets is approximately 40 degrees. The most interesting and novel structural feature lies in the beta-arches connecting the two sheets. The only two beta-arches in B07 are located on the same side of the crossover region, leaving the other side completely unconnected. This special arrangement presents an interesting fact that B07 lacks the “interlock” arrangement, a pattern also referred to as the cross-*β* motif^3^ that in definition requires the two beta-arches formed by adjacent beta strands to interlock with each other (Supplementary Figure S18). Historically, the cross-*β* motif was considered as a general constraint for almost all beta sandwiches^2,4^ and subsequently adopted as a primary design principle for new proteins^3^. Also interestingly, the hairpin loop between the first and second beta strands consists of 10 residues, significantly longer than the usual case, and extends to form hydrophobic interactions with another hairpin loop on the opposing sheet. This special loop-based structure essentially acts as a cap to cover the unconnected crossover region and protect the hydrophobic core, likely playing a crucial role in the stabilization of the structure.

B08 also features an interconnected alpha helix inserted into the interlock region between the two beta sheets. However, unlike B10, B08 has a less compact shape, with the alpha helix being less exposed, which may explain its relatively lower thermostability.

B21, with a total of 200 residues, is the longest design having a separable monomeric state. The two beta sheets in B21 have five and eight strands, respectively, with each strand ranging from 5 to 15 residues. Each plane of the two sheets is highly twisted, forming a compact overall structure.

Overall, the novel structural characteristics in these successful designs offer new insights into the underlying rules governing the design space of mainly beta proteins, opening up new possibilities for their design and application.

